# SlWRKY80 recruits SlHDA1 to regulates the tomato fruit ripening and color transformation

**DOI:** 10.1101/2024.01.31.578225

**Authors:** Chunyu Shang, Guo Chen, Xiaoyan Liu, Hao Zheng, Guobin Li, Jiafa Wang, Songshen Hu, Zhenglun Li, Xiaohui Hu

**Author notes:** Corresponding authors: Xiaohui Hu. E-mail addresses (C. Shang), (G. Chen), (X. Liu), (H. Zheng), (G. Li), (S. Hu), (Z. Li), (X. Hu).

## Abstract

Fruit ripening involves key transcription factors (TF). Despite the established importance of WRKY TFs in plant development, the involvement of WRKY and its underlying mechanism in the regulation of fruit ripening and color transformation remain unclear. Here, we discovered a phenomenon where SlWRKY80 is involved in the ripening and color transformation of tomato. Overexpression of *SlWRKY80* shortened the color transition cycle of tomato fruit, whereas loss-of-function mutation in *SlWRKY80* delays this process. On the 7^th^ day after the color breakdown (Break+7), the fruit color was bright red in *SlWRKY80*-overexpression lines. SlWRKY80 could bind to downstream promoters of *SlSGR1* and *SlPSY1*, thereby exerting a positive regulatory effect on the expression levels of *SlSGR1* and *SlPSY1*. This regulatory activity led to the facilitation of chlorophyll a decomposition and carotenoid synthesis during the transition of fruit color, ultimately accelerating fruit ripening and color change. The *SlWRKY80* knockout lines (*slwrky80^cr^*-3 and *slwrky80^cr^*-4) showed light red. Thr (The 69^th^ amino acid) mutated to Ser of SlWRKY80 (*slwrky80^cr^*-46), SlHDA1 was recruited as a regulatory factor to enhance the regulation of downstream *SlPSY1* by *SlWRKY80*, and inhibited the *SlSGR1* expression by *SlWRKY80* which maybe the reason why the *slwrky80^cr^*-46 knockout lines exhibited brown color. In conclusion, SlWRKY80 regulates the expression of *SlSGR1* and *SlPSY1* by recruiting SlHDA1, thereby regulating chlorophyll decomposition and carotenoid synthesis, and further participating in fruit maturation and related fruit color formation. This research has important guiding significance for breeding tomato germplasm with excellent commercial traits.

## Introduction

Tomato (*Solanum lycopersicum* L.) is a model system to study the mechanism of fruit ripening (Li et al. 2020; Jiang et al., 2023). Tomato fruit ripening includes changes in fruit color, increased flavor compounds, and fruit softening, and so on. Color quality, an important aspect of tomato fruit quality, makes an irreplaceable contribution to the sensory and nutritional quality of the fruit (Ilahy et al., 2018). During the ripening process of tomatoes from green to red, chlorophyll continuously degrades, whereas lycopene, β-carotenoids and other colorants constantly accumulate (Lin et al., 2014). The ripening cycle of tomato is not only closely related to fruit quality but also to shelf life and market prices. Therefore, exploring the internal mechanisms of tomato fruit ripening and color transformation has great guiding significance in selecting tomato germplasm with excellent commercial characteristics (Li et al., 2019).

Numerous studies have reported transcription factors (TFs) involved in the regulation of ripening. To comprehend the process of fruit ripening regulation, it is crucial to investigate the connections between higher level regulatory genes, intermediate regulators and downstream effectors. WRKY TFs are one of the largest specific transcription factor families in plants. They involve in plant growth and development, response to abiotic and biotic stresses, play a dynamic role in secondary metabolic regulation, and plant hormone signaling processes (Zhang et al., 2022). Increasing numbers of SlWRKY TFs have been characterized by fruit ripening regulators and color transformation in tomato. *SlWRKY16/17/53/54* gene exhibits a strong association with the physiological process of tomato ripening, with particular emphasis on the involvement and significance of *SlWRKY17/53* in the regulatory mechanisms governing ripening (Liu et al., 2022). SlWRKY TFs may engage in the modulation of tomato fruit ripening and senescence through diverse pathways, including but not limited to chlorophyll degradation, lycopene synthesis, and the participation of transcription factors ERF or RIN (MADS-box transcription factor ripening inhibitors) that are closely linked to the ripening and senescence of fruits (Liu et al., 2022). RIN also regulates the accumulation of carotenoids by interacting with the *SlPSY1* promoter (Martel et al., 2011). The STAY-GREEN 1 (*SlSGR1*) gene is a regulatory target of RIN. *SlSGR1* engages in direct interaction with *SlPSY1* during fruit ripening, resulting in the inhibition of *SlPSY1* activity. This inhibition subsequently leads to the accumulation of lycopene and β-carotenoids (Fujisawa et al., 2013; Hortenstein, 2009; Luo et al., 2013). The regulation of *SlWRKY35* in mature tomato mediated by the regulatory factor SIRIN, thereby directly stimulating the expression of *SlDXS1*. This activation leading to an enhanced accumulation of carotenoids. Furthermore, the co-overexpression of *SlWRKY35* and *SlLCYE* in tomato fruits can augment lutein production, expedite tomato ripening, and facilitate color transformation (Yuan et al., 2022). The involvement of *SlWRKY22* in the coordinated regulation of the expression levels of six calcium-sensing genes is instrumental in the regulation of tomato fruit ripening (Arhondakis et al., 2016). These findings suggest that the WRKY TFs actively contributes to the ripening process of tomato fruits and assumes a crucial function in the modulation of fruit color changes in conjunction with other regulatory factors (Wang et al., 2017).

The hue of the fruit depends on the pigmentation of the outer epidermis and inner flesh. Chlorophyll degradation is considered the initial step in color change during fruit ripening. Previous studies that the *SlSGR1* mutation (*gf*) mutant tomato still retains chlorophyll during ripening (Akhtar et al., 1999). The gene that causes the green retention phenotype of the *gf* mutant is named *SlSGR1* (Barry et al., 2008). Currently, the research on the green retention phenotype of the *SlSGR1* mutant is extensive in the leaves, and the withered leaves of the mutant ultimately metabolize but remain green. The literature regarding the underlying mechanism of abnormal browning in the fruit of the tomato mutant *SlSGR1* is limited (Luo et al., 2013; Wang et al., 2022).

The accumulation of lycopene is a distinctive phenomenon observed during the ripening of tomato fruits. Studies revealed that the *SlPSY1* enzyme facilitates the condensation of geranylgeranylgeranyl pyrophosphate (GGPP) to generate phytoene. This serves as the initial step in the carotenoid synthesis pathway and acts as a crucial rate-limiting step in carotenoid production. Mutations in this gene can impede lycopene synthesis and the development of red tomato fruits (Kachanovsky et al., 2012). The gene *SlPSY1* plays a crucial role in regulating phytoene synthesis (Summers et al., 1993; Wang et al., 2021; Xiong et al., 2019; Wu et al., 2020). The association between SlWRKY TFs and *SlPSY1* is highly significant. SlWRKY16/17/3 individually binds to the *SlPSY1* promoter, thereby governing the production of phytoene. SlWRKY16 and SlWRKY17 exhibit interaction at the protein level, leading to a substantial enhancement in the regulation of the *SlPSY1* promoter (Wang et al., 2017).

Deacetylation modifications mediated by histone deacetylases (HDACs) play an important role in tomato fruit ripening and color transformation processes (Guo et al., 2017). *SlHDA1* can interact with *TAG1* or *TM29*, which are associated with fruit ripening. Conversely, *SlHDA3* exclusively interacts with *TAG1*, thereby governing fruit development, ripening, and aging (Zhao et al., 2015). Guo et al. (2017, 2018, 2022) demonstrated that *SlHDA1*, *SlHDA3*, and *SlHDT1* play a negative role in regulating tomato fruit ripening and carotenoid accumulation and suppressing these genes considerably expedites the ripening process of tomato fruits. However, research investigating the involvement of DHACs interact with WRKY proteins to tomato fruit ripening is scarce (Kim et al.,2008).

In this study, we comprehensively characterize the roles of *SlWRKY80* in regulating tomato ripening and color transition process by carotenoid metabolism. *SlWRKY80* knockout delayed tomato ripening and color transformation, and different knockout types also had a significant impact on fruit color. This study conducted in-depth research on the mechanism of the emergence of special phenotypes in tomato fruits, and reveals a novel *SlWRKY80*-dependent regulatory mechanism that involves carotenoid metabolism pathway between SlWRKY80 and SlHDA1 to regulate tomato fruit ripening and color transformation. Our results unravel a previous undescribed regulatory module of fruit ripening in tomato.

## Result

### Overexpression of *SlWRKY80* shortens tomato color transition cycle

To gain insight into the possible involvement of *SlWRKY80* in the regulation of fruit ripening, we successfully generated overexpressed *SlWRKY80* lines (80OE-1 and 80OE-3) and *SlWRKY80* knockout mutants (*slwrky80^cr^*-3, *slwrky80^cr^*-4, and *slwrky80^cr^*-46) via genetic transformation. Phenotypic analysis showed that overexpressing *SlWRKY80* accelerated the fruit maturation process, as evidenced by a reduction of approximately 4 days in the duration from flowering to fruit color breakdown (Supplementary table 1), whereas *SlWRKY80* knockout lines fruits displayed a delayed fruit ripening phenotype (average 3d late). Moreover, the overexpressing *SlWRKY80* shortened the fruit color transition cycle after the color breakdown period (break), whereas the *SlWRKY80* knockout lines (*slwrky80^cr^*-3 and *slwrky80^cr^*-4) showed the opposite trend. Additionally, the fruit of *slwrky80^cr^*-46 line manifested a brown phenotype during color transition, thereby complicating the findings further (Fig. 1A). No notable alterations were observed in the levels of chlorophyll a (Fig. 1B) and carotenoids (Fig. 1D), fruit firmness (Fig. 1F), and soluble solid content in fruit (Fig. 1G) throughout the color breaking phase of all the transgenic lines. However, compared with the other knockout lines, the *slwrky80^cr^*-46 lines significantly elevated chlorophyll b content (Fig. 1C).

**Figure 1.**
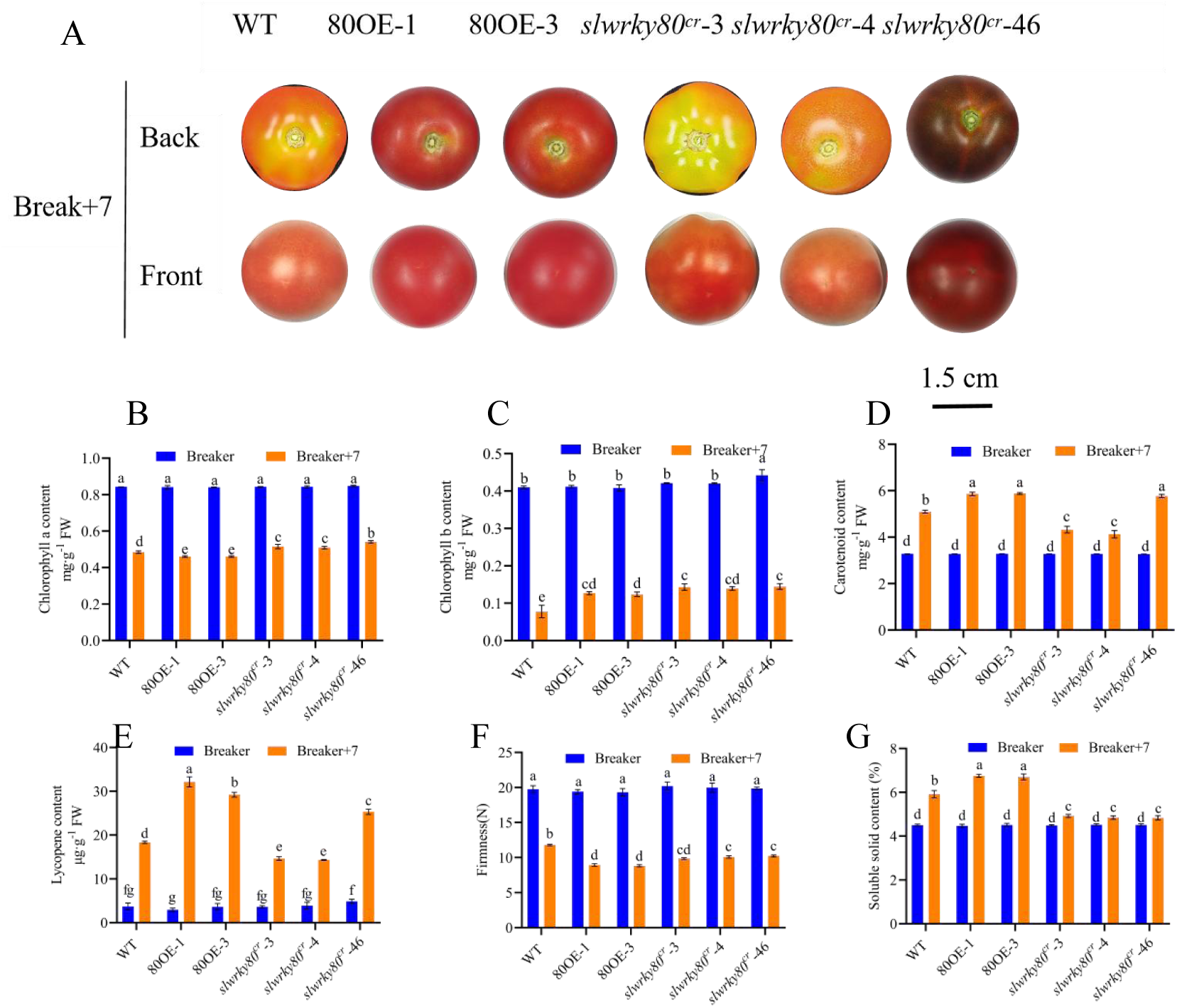
Overexpressing *SlWRKY80* shortens the color transition cycle by promoting chlorophyll a degradation and carotenoid synthesis of tomato fruit in different lines during the Break+7 period. A, Phenotypic maps of ripening fruit. B, Chlorophyll a content. C, Chlorophyll b content. D, Carotenoids content. E, Lycopene content. F, Fruit firmness. G, Soluble solids content. Values represented in as the mean ± SD of 3 biological replicates. Different letters above the bars indicate statistically significant differences between the samples (LSD’s test; *P* < 0.01).

On the 7^th^ day after the color breakdown period (Break+7 period), the chlorophyll a content of the *SlWRKY80* overexpressing lines was significantly lower than that of the WT, whereas the contents of carotenoid and lycopene in the former were significantly higher than those of the latter (Fig. 1D, 1E). Moreover, the *slwrky80^cr^*-3 knockout lines and *slwrky80^cr^*-4 knockout lines were the opposite. However, the *slwrky80^cr^*-46 lines with a brown phenotype had the highest chlorophyll a content and were significantly higher than other knockout lines. The carotenoid contents in *slwrky80^cr^*-46 knockout lines and the *SlWRKY80* overexpressing lines had no significant difference, and they were significantly higher than those of the WT, *slwrky80^cr^*-3, and *slwrky80^cr^*-4 knockout lines. Furthermore, compared with the lycopene content in *SlWRKY80* overexpressing lines, that of *slwrky80^cr^*-46 lines exhibited a statistically significant decrease during the Break+7 period. The lycopene content in knockout *slwrky80^cr^*-46 lines displayed a statistically significant increase compared to the lycopene contents of the WT, *slwrky80^cr^*-3, and *slwrky80^cr^*-4 lines. However, compared with the *slwrky80^cr^*-3 and *slwrky80^cr^*-4 lines, the *slwrky80^cr^*-46 lines had no statistically significant differences in fruit firmness (Fig. 1F) and soluble solid content (Fig. 1G) in the fruit.

In summary, the upregulation of *SlWRKY80* expedites the breakdown of chlorophyll a, thereby facilitating premature fruit maturation and concurrently augmenting the levels of carotenoids and soluble solids postcolor transition. Conversely, the *SlWRKY80* knockout lines (*slwrky80^cr^*-3 and *slwrky80^cr^*-4) exhibit contrasting outcomes. Furthermore, the observed fruit browning in the *slwrky80^cr^*-46 lines may be correlated with the significant increase in carotenoids and a significant decrease in chlorophyll a decomposition.

### SlWRKY80 binds downstream promoters of *SlSGR1* and *SlPSY1* and positively regulates their expression levels

On the basis of existing reports, the genes related to chlorophyll metabolism and lycopene synthesis, including *SlSGR1*, *SlPSY1*, *SlPAO*, *SlPPH*, and *SlNYC1* were identified (Luo., 2013; Guo et al., 2017; Ding et al., 2022). All of these genes were upregulated in the overexpressing *SlWRKY80* lines (Fig. 2A). Among the stages of Break, Break+7, and Break+10, the relative expression levels of *SlSGR1* and *SlPSY1* most significantly changes, and the overexpressing *SlWRKY80* significantly increased the relative expression levels of *SlSGR1* and *SlPSY1*. Knocking out *SlWRKY80* significantly inhibited the relative expression levels of *SlSGR1* and *SlPSY1* (Figs. 2B and 2C).

**Figure 2.**
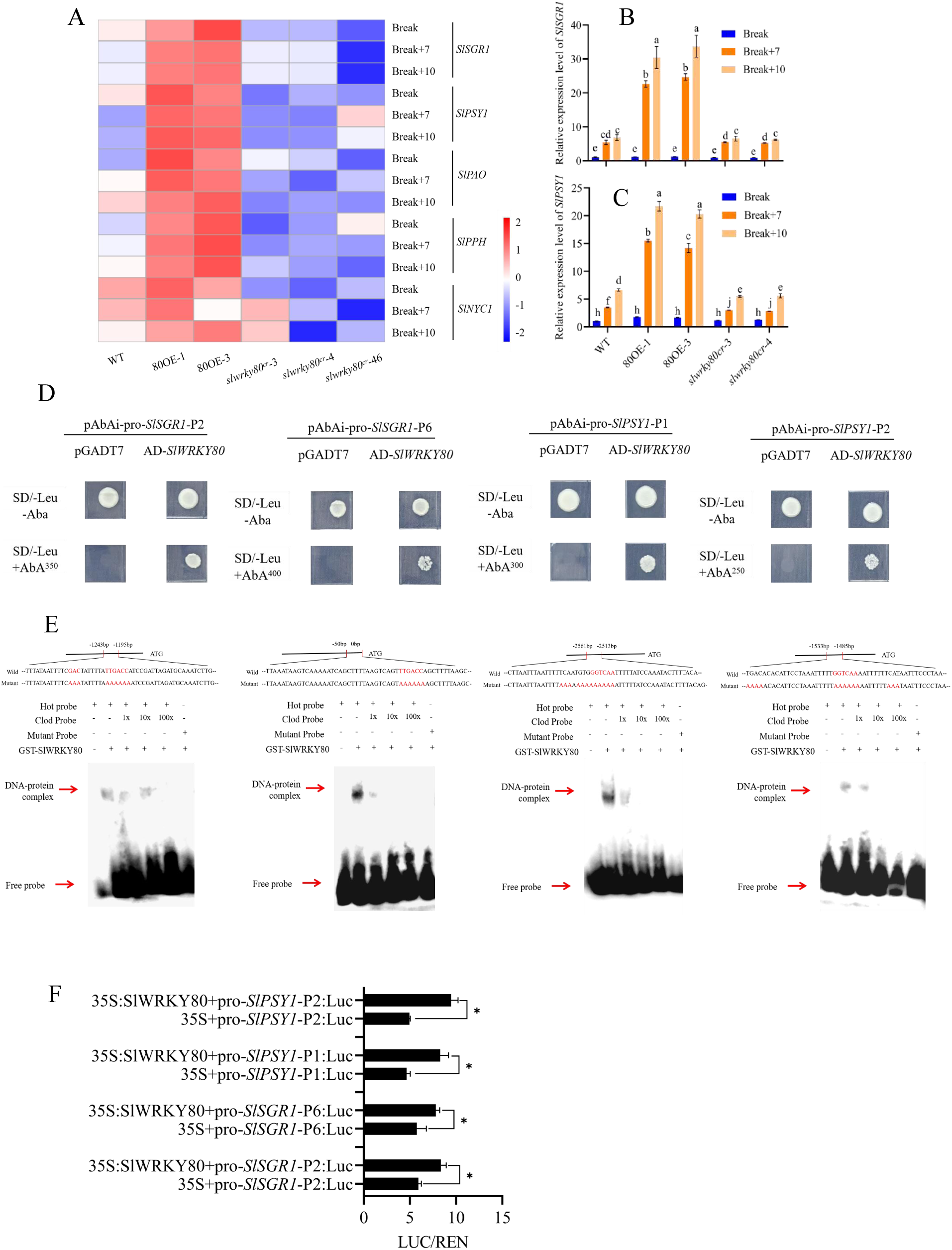
SlWRKY80 binds downstream promoters of *SlSGR1* and *SlPSY1* and positively regulates their expression. A, Heat map of gene expression of *SlSGR1* and other genes in the fruits of different lines during the Break+10 period. B and C, Relative expression levels of *SlSGR1* and *SlPSY1* in fruits of different lines during the Break+10 period, respectively. D and E, Y1H and EMSA validation of SlWRKY80 binding to *SlSGR1* and *SlPSY1* promoters in vitro. F. D-Luc validation of SlWRKY80 positively regulates the promoters of *SlSGR1* and *SlPSY1* in vivo. Values represent the mean s the mean ± SD of 3 biological replicates, significant differences (*P*<0.05) are indicated by different lowercase letters and * based on LSD’s test.

The upstream 3000 bp promoter region of *SlSGR1* contains six W-boxes, whereas that of *SlPSY1* contains two W-boxes (Supplement fight 1). Subsequently, Y1H (Fig. 2D) and EMSA (Fig. 2E) revealed the binding capability of SlWRKY80 to the promoters of *SlSGR1* (P2, P6) and *SlPSY1* (P1, P2). The experiments in vivo provided evidence of SlWRKY80’s positive regulation of the expression levels of *SlSGR1* and *SlPSY1*. SlWRKY80 participates in the process of tomato fruit color ripening and color transformation by regulating *SlSGR1* and *SlPSY1*.

### The *slwrky80^cr^*-46 lines exhibits a brown fruit phenotype

Transmission electron microscopy (TEM) provides a comprehensive and intuitive means of examining the microscopic composition of tomato fruit from the microscopic level (Luo et al., 2013). In this experiment, WT, *slwrky80^cr^*-4, and *slwrky80^cr^*-46 pulp were selected for TEM (Fig. 3A), and samples were taken at Break and Break+7 periods. TEM analysis revealed that the chloroplast undamaged, and the thylakoid grana layer was distinctly discernible in fruits of all lines during the Break stage. At Breaker+7 stage, the thylakoid grana layer in WT and *slwrky80^cr^*-46 exhibited signs of decomposition, whereas the number of plastid spherules increase, resulting in the lycopene crystals accumulation significantly. Conversely, *slwrky80^cr^*-4 chloroplasts maintained a relatively intact thylakoid grana layer, with no discernible presence of lycopene crystals. By contrast, the thylakoid grana decomposition and the lycopene crystal accumulation were accelerated and the number of plastid globules increased in *slwrky80^cr^*-46 line fruit (Fig. 3B). These cytoplasmic globular structures and lycopene crystals play a crucial role in the synthesis and storage of pigments, including lycopene. Therefore, the brown phenotype observed in *slwrky80^cr^*-46 fruit related to the rate of chlorophyll decomposition and the synthesis of chromogenic pigments such as lycopene.

**Figure 3.**
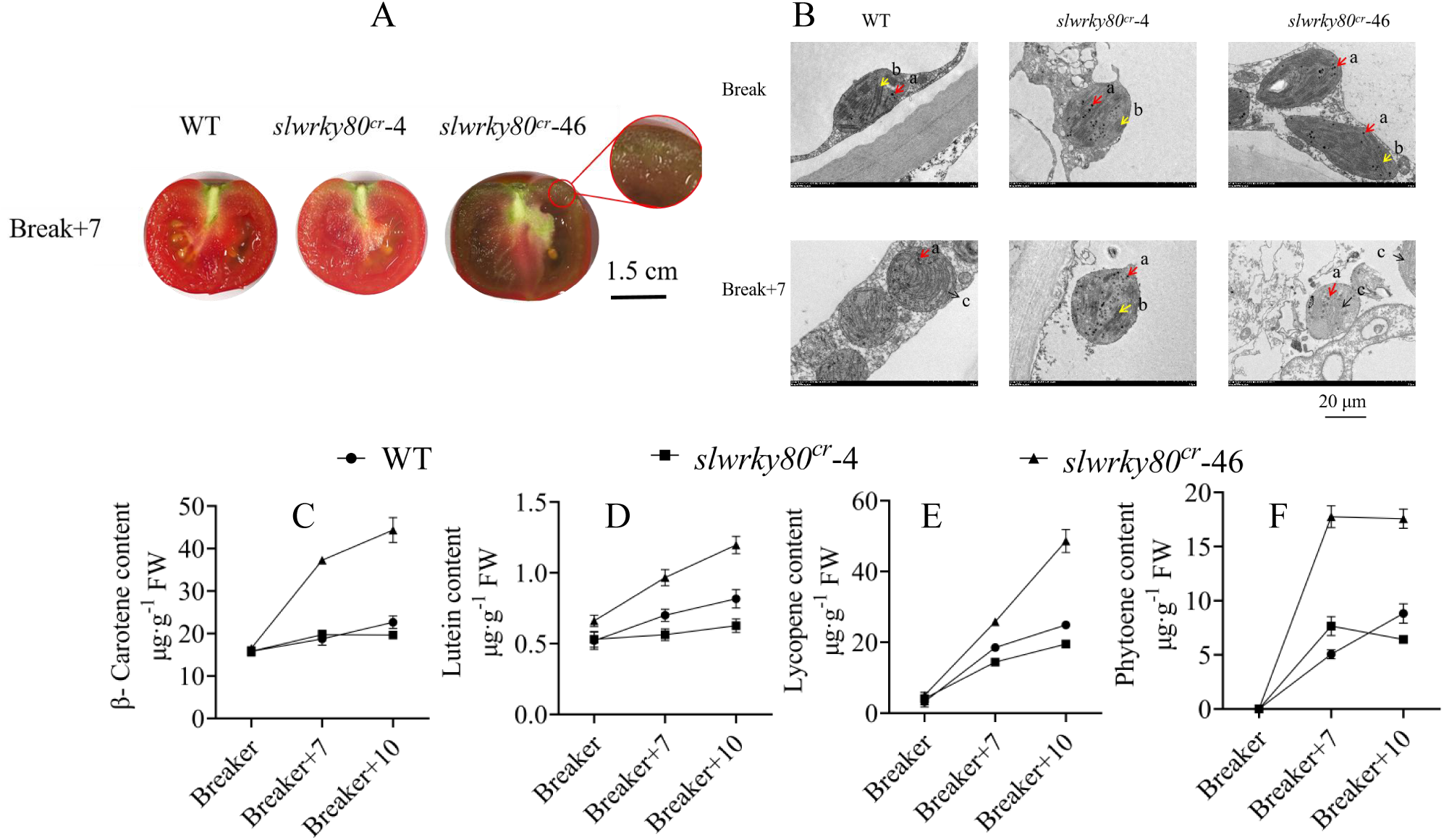
*slwrky80^cr^*-46 knockout lines exhibits a fruit brown phenotype. A, Cross section diagram of WT, *slwrky80^cr^*-4, and *slwrky80^cr^*-46 fruits during Break+7 period. The red circle represents the sampling site for TEM. B, TEM images of fruits during Break and Break+7 periods. Note: a, The red arrow indicates placeglobles. b, The yellow arrow indicates thylakoid grana. c, The black arrow represents the lycopene crystal. C-F, The contents of β-carotene, lutein, lycopene, and phytoene in WT, *slwrky80^cr^*-4, and *slwrky80^cr^*-46 fruits during Break, Break+7, and Break+10 periods. Values represent the mean across three biological replicates, error bars represent SD, significant differences (*P*<0.05) are indicated by lowercase letters based on LSD’s test.

Carotenoid metabolism is involved in the whole fruit color conversion cycle. So we determined 66 substances in the carotenoid metabolism pathways of WT, *slwrky80^cr^*-4, and *slwrky80^cr^*-46 lines (Supplementary figure 2). We also compared the GO classification of WT, *slwrky80^cr^*-4, and *slwrky80^cr^*-46 during Break+7 periods (Supplementary figure 3) to investigate further the reason for the browning of the fruit of *slwrky80^cr^*-46. At the same time, we analyzed the gene variation volcano maps (Supplementary Figure 5), GO classification maps (Supplementary Figure 6), KEGG metabolism (Supplementary Figure 7), and the DEG gene counts (Supplementary Figure 4) of *slwrky80^cr^*-4 and *slwrky80^cr^*-46 lines compared to WT during Break and Break+7 periods. These data all indicate a significant correlation between the brownish color and carotenoids in the fruit of *slwrky80^cr^*-46 lines.

The levels of β-carotene (Fig. 3C), lutein (Fig. 3D), lycopene (Fig. 3E), and phytoene (Fig. 3F) in the *slwrky80^cr^*-46 lines were the highest, thus we infer that the abnormal increase in carotenoid content is responsible for the brown fruit observed in the *slwrky80^cr^*-46 lines.

### Amino acid mutated from Thr to Ser lead to brown fruit of *slwrky80^cr^*-46

We analyzed three gene editing types and the amino acid composition of the *SlWRKY80* knockout lines to investigate the factors contributing to the manifestation of the brown phenotype in the fruit of *slwrky80^cr^*-46. We found that a substitution of Thr with Ser at the 69^th^ amino acid position in *slwrky80^cr^*-46 (Fig. 4A). This substitution was accompanied by notable alterations at position 69 and neighboring phosphorylation sites (Fig. 4B, Supplementary table 2).

**Figure 4.**
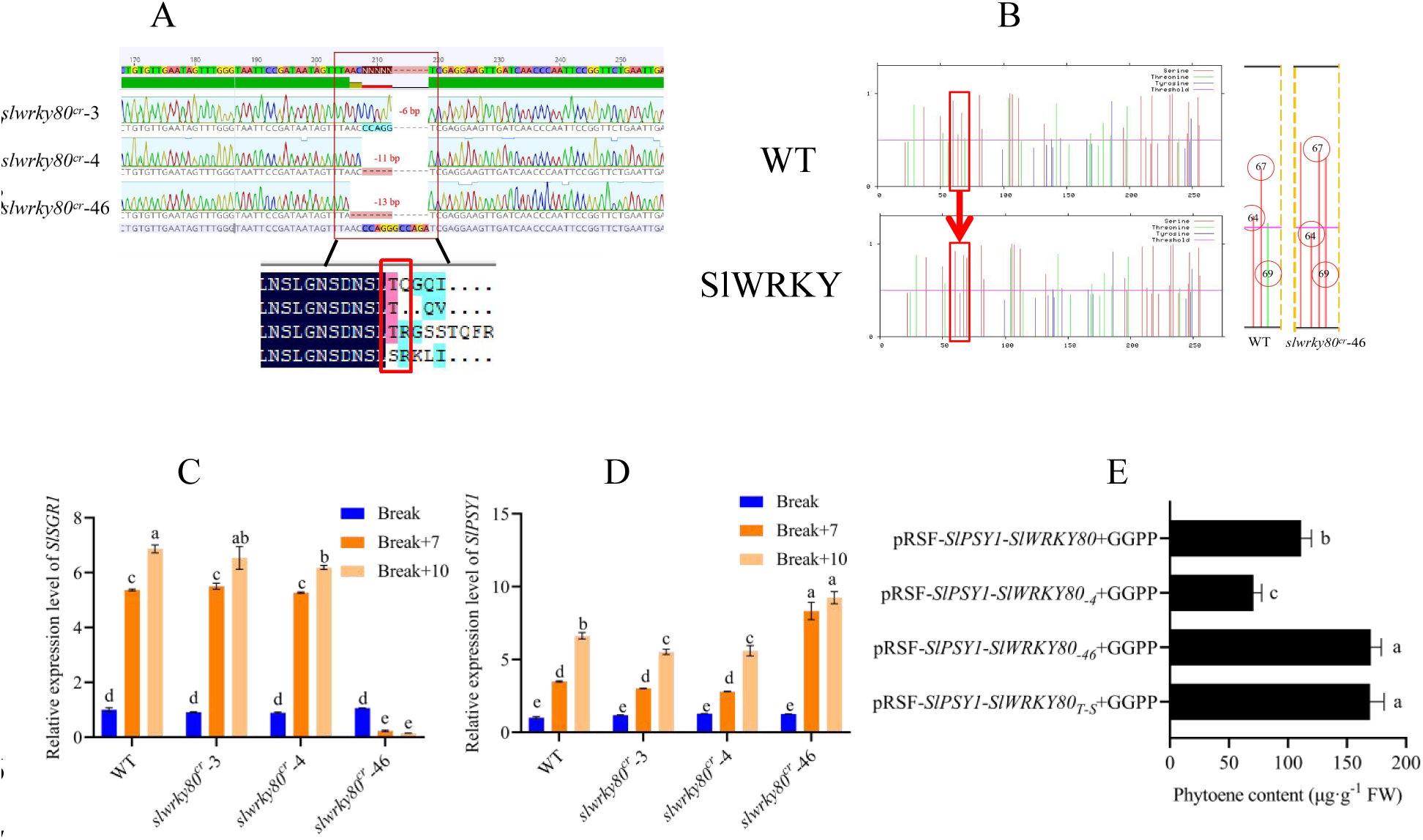
The brown color and amino acid Thr mutation to Ser in the tomato fruits of the *slwrky80^cr^*-46 lines are related. A. SlWRKY80 knockout lines and DNA and amino acid editing types. B. Analysis of phosphorylation sites in WT and SlWRKY80-T-S. C and D. The relative expression levels of *SlSGR1* and *SlPSY1* in WT, *slwrky80^cr^*-4, and *slwrky80^cr^*-46 at Break, Break+7, and Break+10 periods. E. Engineering bacterial experiments verified that SlWRKY80-T-S can significantly increase the content of phytoene. Values represent the mean across three biological replicates, error bars represent SD, and significant differences (*P*<0.05) are indicated by lowercase letters based on LSD’s test.

The relative expression trends of *SlSGR1* and *SlPSY1* remained consistent across the three stages of Break, Break+7, and Break+10 among three *SlWRKY80* knockout lines. Relative expression levels of *SlSGR1* and *SlPSY1* were not significantly altered in *slwrky80^cr^*-3 and *slwrky80^cr^*-4 were observed at Break+7. Relative expression levels of *SlSGR1* and *SlPSY1* were significantly higher than WT at Break+7 and Break+10, whereas *slwrky80^cr^*-46 was significantly lower than WT (Fig. 4C). At Break, Break+7, and Break+10 stages, the expression levels of *SlPSY1* in *slwrky80^cr^*-3, *slwrky80^cr^*-4 and *slwrky80^cr^*-46 significantly increase. The relative expression levels of *SlPSY1* in the *slwrky80^cr^*-46 lines significantly higher than those observed in the other lines (Fig. 4D).

To further verify the relationship between the mutation of SIWRKY80 amino acid from Thr to Ser and the synthesis of phytoene, we validated it through engineering bacterial experiments. After transferring the pRSF-*SlPSY1*-*SlWRKY80* (−4/-46/-T-S) fusion vector capable of prokaryotic expression of *SlPSY1* into engineering lines carrying a GGPP substrate for synthesizing phytoene, the expression of phytoene by the engineering lines showed a significant decrease in pRSF-*SlPSY1*-*SlWRKY80*-4 and a significant increase in pRSF-*SlPSY1*-*SlWRKY80*-46 and pRSF-*SlPSY1*-*SlWRKY80*-T-S (Fig. 4E). Taken together, the mutation of SlWRKY80 amino acid from Thr to Ser affects the mutation site and adjacent phosphorylation sites, significantly inhibiting the expression of *SlSGR1*. Moreover, it specifically increases the expression of phytoene, leading to an abnormal increase in phytoene content.

### SlHDA1 was recruited by SlWRKY80 to participate in the appearance of brown fruits

Thus far, the reason for the presence of brown fruit in *slwrky80^cr^*-46 has remained unclear. We combined existing reports indicating that the SlHDA family participates in tomato fruit ripening by affecting the accumulation of carotenoids to understand deeply how the amino acid mutation from Thr to Ser in SlWRKY80 leads to the manifestation of brown color in the tomato fruit (Guo et al., 2017; Guo et al., 2018; Guo et al., 2022). The heatmaps of 13 related genes in the SlHDA family were examined in the SlWRKY80 transgenic lines (Fig. 5A). Except for *SlHDA4*, the *slwrky80^cr^*-3 lines and *slwrky80^cr^*-4 lines exhibited an upregulation trend. Conversely, the expression of *SlHDA1/3/4/6/8/10* in the *slwrky80^cr^*-46 lines displayed a downregulation trend. The protein level interaction between SlWRKY80 and SlHDA1/3/9 were validated via in vitro experiments Y2H (Fig. 5B). The Pull-down test further confirmed the protein level interaction between SlWRKY80 and SlHDA1 (Fig. 5C). However, the Pull-down test could not verify the interaction between SlWRKY80 and SlHDA3 and SlHDA9. The protein structure of SlWRKY80 was predicted by utilizing Alphafold2 prior to and subsequent to a single amino acid (T-S) mutation, along with its interaction with SlHDA1. The analysis revealed that the T-S mutation induced substantial alterations in the protein structure of SlWRKY80, as well as a notable modification in its binding affinity towards SlHDA1 (Fig. 5D). We predicted the binding affinity and dissociation constant at 25°C of SlWRKY80 before and after single amino acid (T-S) mutation with Prodigy and found that the binding affinity and dissociation constant at 25°C were high after the T-S mutation. This finding suggests that the interaction strength between SlWRKY80 and SlHDA1 significantly decreased after the T-S mutation. At the Break+7 stage, the overexpressing *SlWRKY80* significantly inhibited the expression of *SlHDA1*, whereas the *slwrky80^cr^*-3 lines and *slwrky80^cr^*-4 lines had the opposite effect. However, the expression level of *SlHDA1* in *slwrky80^cr^*-46 lines was further reduced (Fig. 5E). We conducted a D-Luc assay to demonstrate further the significance of this weakened interaction. We found that after the T-S mutation in the amino acid of SlWRKY80 (*slwrky80^cr^*-46 and *slwrky80^cr^*-T-S), the regulation of the *SlSGR1* promoter in the presence of SlHDA1 was further weakened. However, the regulation of the *SlPSY1* promoter was significantly increased (Fig. 5F). Above all, SlHDA1 was recruited by SlWRKY80 to participate in the maturation of tomato fruits and the appearance of brown fruits.

**Figure 5.**
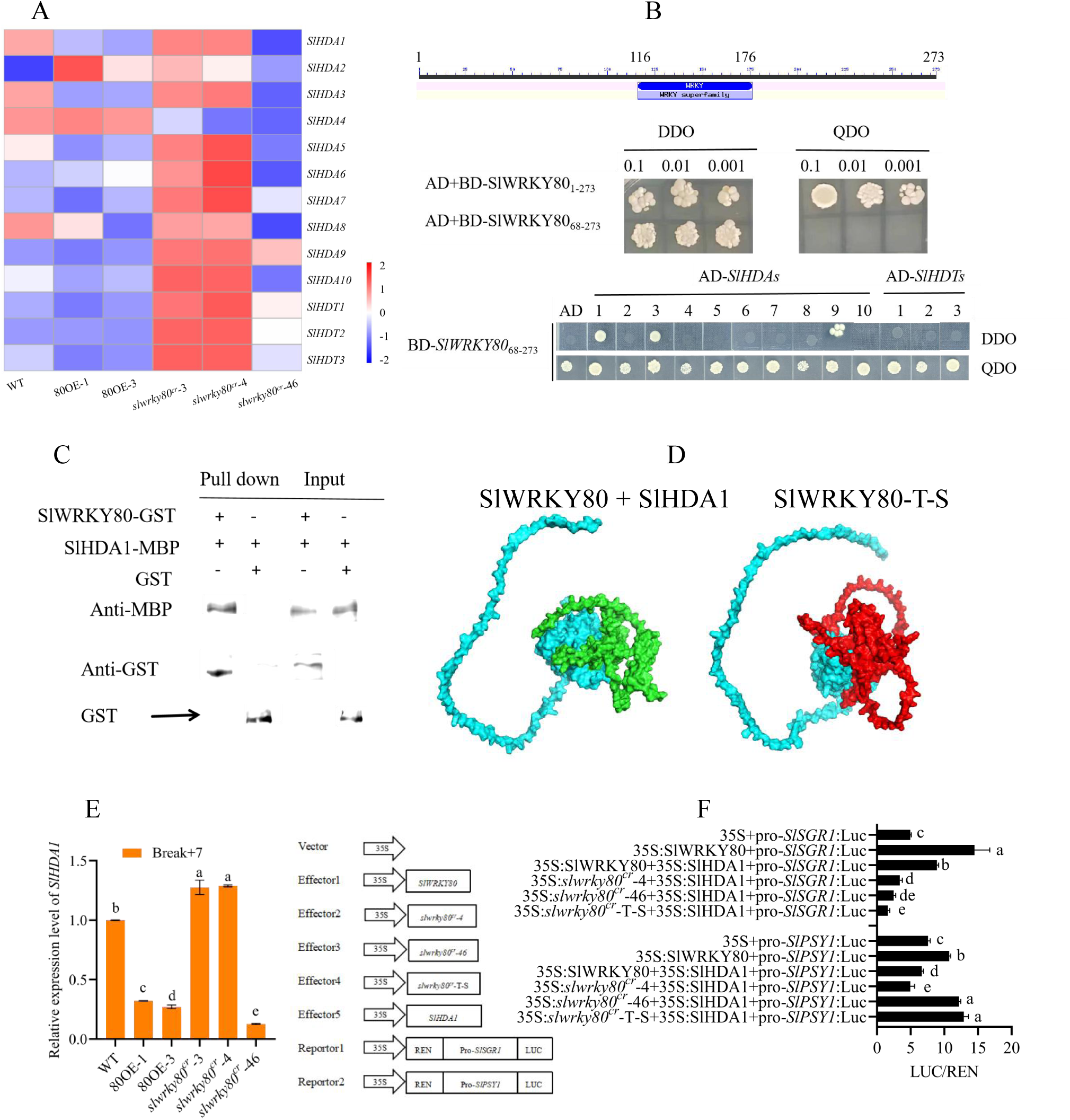
SlWRKY80 recruited SlHDA1 to participate in tomato fruit ripening and color transformation. A, Heat maps of SlHDAs gene expression in the fruits of each lines at Break, Break+7, and Break+10 stages. B and C, Verification of the interaction between SlWRKY80 and SlHDAs proteins through Y2H and Pull-down experiments. D, Prediction of the structure of SlWRKY80 amino acid mutation (Thr-Ser) and SlHDA1 protein interaction using Alphafold2. A:Green represents SlWRKY80 protein. B: Blue represents SlHDA1 protein. C: Red represents SlWRKY80-T-S protein. E, The relative expression levels of SlHDA1 in each lines during Break+7 period. F, The D-Luc experiment verified that SlHDA1 plays an important regulatory role in the ripening and color transformation process of SlWRKY80 fruit. Values represent the mean across three biological replicates, error bars represent SD, significant differences (*P*<0.05) are indicated by lowercase letters based on LSD’s test.

**Figure 6.**
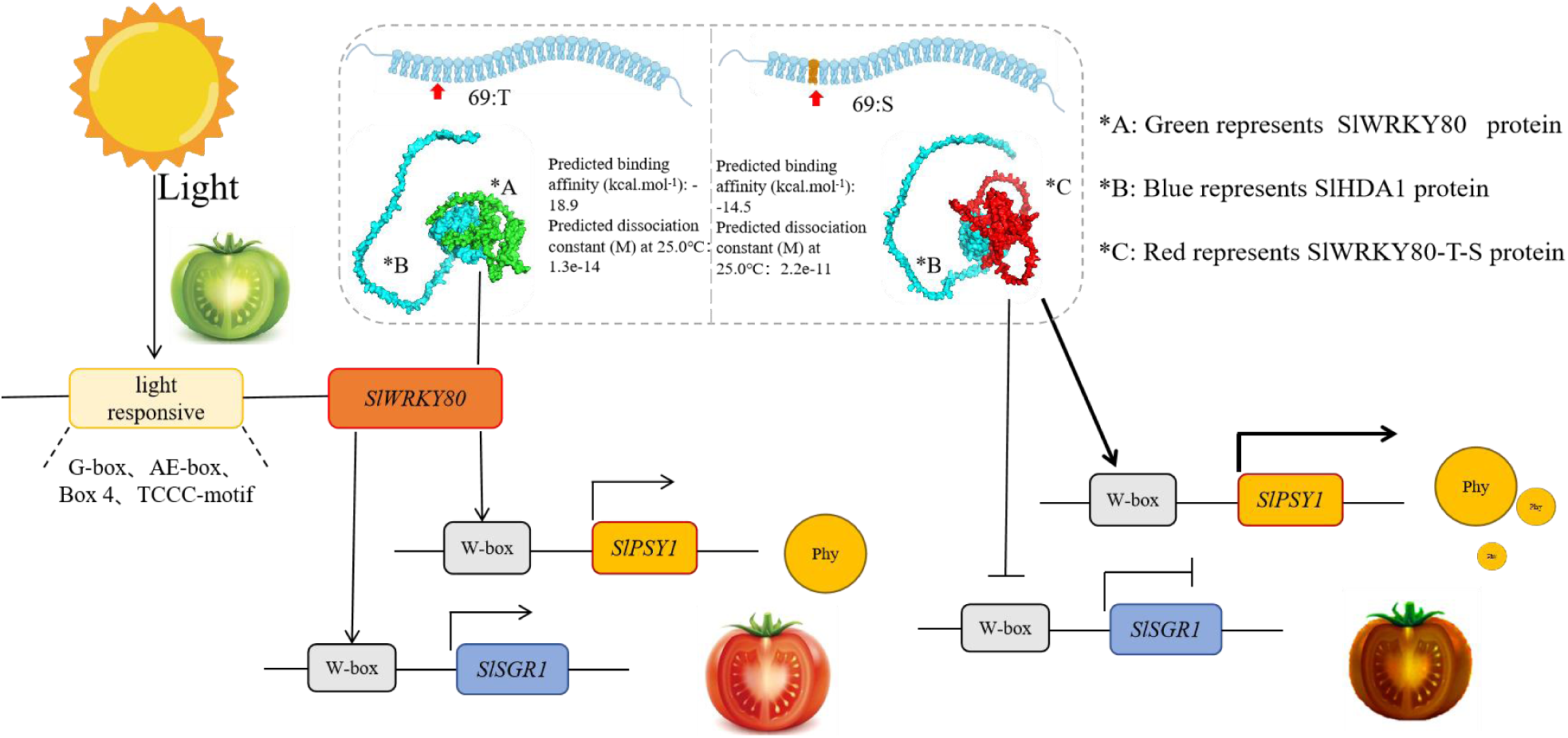
Model of SlWRKY80 recruiting SlHDA1 to participate in fruit ripening and color transformation.

## Discussion

### Overexpression of *SlWRKY80* shortening tomato color transition cycle

The ripening and color change of tomato fruit are inseparable from the carotenoid metabolic pathway. Carotenoids are a class of natural organic pigments that are widely found in plants. Carotenoids consist of two main categories: lutein and carotene, which include lycopene, neurosporene, α-carotene, β-carotene, and γ-carotene, among others. Lycopene is currently the strongest antioxidant found in nature (Sun et al., 2022). During the ripening process of tomato fruits from green to red, chlorophyll degrades, and lycopene, beta-carotene, lutein accumulate gradually which is accompanied by the softening of the fruit and other changes. As one of the largest families of transcription factors in plants, WRKY also plays a significant role in regulating fruit ripening and color change. *SlWRKY16/17/22/35/ 53/54* are involved in fruit ripening and color change through carotenoid metabolism-related pathways (Liu et al., 2022).

In this study, overexpressing *SlWRKY80* shortens the fruit ripening cycle, with an average reduction of 4 days from the flowering stage to the onset of fruit color break (Supplementary table 1). Moreover, the overexpressing *SlWRKY80* reduces the tomato color change period, resulting in a bright red fruit color on the 7^th^ day after the color break. By contrast, the *SlWRKY80* knockout lines (*slwrky80^cr^*-3 and *slwrky80^cr^*-4) exhibit the opposite effect. Another interesting discovery is that the *slwrky80^cr^*-46 line shows a brown fruit phenotype, which is different from that shown by other knockout types (Fig. 1A). Overexpression of *SlWRKY80* accelerates the degradation of chlorophyll a to advance fruit maturation, and increase carotenoid and soluble solids contentat the Break to Break+7 period (Fig. 1). Conversely, the *slwrky80^cr^*-3 and *slwrky80^cr^*-4 knockout lines show the opposite effects. The contents of carotenoid and lycopene are significant increase which may be related to the brown color of the *slwrky80^cr^*-46 lines fruits.

### SlWRKY80 binds downstream *SlSGR1* and *SlPSY1* promoters to positively regulate the expression of *SlSGR1* and *SlPSY1*

Wang et al. (2017) screened 23 differential SlWRKY transcription factors, including SlWRKY80, from the mature transcripts of tomato fruits, but the relative expression level of *SlRWKY80* was not significant during tomato ripening. This discrepancy can potentially be attributed to variations in tomato species and maturity stages employed during the experiment. Wang et al. (2017) demonstrated that SlWRKY16/17/3 can bind to the *SlPSY1* promoter, thereby exerting regulatory control over the synthesis of phytoene. Additionally, the interaction between SlWRKY16 and SlWRKY17 occurs at the protein level, and their co-overexpression leads to a substantial enhancement in the regulation of the *SlPSY1* promoter. Chlorophyll degradation occurs through the involvement of PAO and chlorophyll oxidation bleaching pathways, whereas SGR encodes a demethylase responsible for catalyzing the degradation of Chla. Suppression or inhibition of SGR expression results in a delay in Chla degradation, leading to the manifestation of a green hysteresis phenotype in plants during natural development and dark-induced aging processes (Zhang et al., 2022).

In this study, the overexpression of *SlWRKY80* can accelerate chlorophyll a degradation. Therefore, we chose *SlSGR1*, which is specifically expressed in fruits, as the research object. The study of *SlSGR1* on tomato fruits is mostly related to the green retention phenotype (Luo et al., 2013). Silencing *SlSGR1* leads to the appearance of a green retention phenotype in tomato fruits, and *SlSGR1* negatively regulates the expression of *SlPSY1*, which is involved in tomato fruit ripening and color transformation (Luo et al., 2013). The relationship between the *SlWRKY* and *SlSGR1* in chlorophyll degradation was also reported. SlWRKY37 interacts with the protein SlVQ7 containing the VQ motif; combined with the W-box in the *SlSGR1* promoter, SlWRKY37 also participates in the yellowing process of tomato leaves (Wang et al., 2022).

We have combined existing reports and verified the mechanism of *SlWRKY80* regulating tomato fruit ripening and color change through carotenoid metabolism through research (Luo., 2013; Guo et al.,2017). The relative expression levels of *SlSGR1* and *SlPSY1* exhibited the most significant variations throughout the entire color transition cycle (Figs. 2B and 2C). *SlSGR1* and *SlPSY1* promoters contain W-box binding elements (Supplement fight 1). Y1H, EMSA and D-Luc in vitro and in vivo confirmed that SlWRKY80 positively regulates the promoters of *SlSGR1* (P2, P6) and *SlPSY1* (P1, P2) (Fig. 2D, 2E, 2F). Therefore, we propose that the overexpression of *SlWRKY80* promotes the expression of these genes by binding to their promoters. Thus, the degradation of chlorophyll a and the synthesis of lycopene are accelerated, ultimately shortening the color transition cycle of tomato.

### SlHDA1 is recruited by SlWRKY80 as a regulatory factor to participate in fruit ripening and brown fruit production

In this study, a unique knockout line (*slwrky80^cr^*-46) exhibits a fruit color similar to that of the *SlSGR1* silent line, which displays brown fruits. This phenotype piqued our intense research interest. The comparative analysis of the gene sequence and amino acid sequence of the central knockout lines indicates that the 69^th^ amino acid Thr in the *slwrky80^cr^*-46 lines mutates to Ser (Fig. 4A). The q-PCR results demonstrate that the relative expression level of *SlSGR1* in the *slwrky80^cr^*-46 lines is significantly lower than that of other knockdown lines. However, the relative expression level of *SlPSY1* in the *slwrky80^cr^*-46 lines is significantly higher than that of other knockdown lines. We found reveal notable alterations at the 69^th^ amino acid position and the neighboring phosphorylation sites subsequent to the Thr mutation to Ser (Fig. 4B, Supplementary table 2). Moreover, the conducted engineering bacterial experiments substantiated that the substitution of Thr with Ser at the 69^th^ of SlWRKY80 amino acid substantially amplifies the concentration of phytoene (Fig. 4E). In conclusion, the alteration of Thr to Ser within the amino acid sequence of *SlWRKY80* plays a pivotal role in the manifestation of brown fruit in the *slwrky80^cr^*-46 tomato line.

There have been reports indicating that the SlHDA family participates in tomato fruit ripening by influencing carotenoid accumulation (Guo et al., 2017; Guo et al., 2018; Guo et al., 2022), which is highly consistent with the work of SlWRKY80 in fruit ripening through the carotenoid pathway in this study. In the investigation of tomato fruit ripening and color transition, the genes *SlHDA1* (Guo et al., 2017), *SlHDA3* (Guo et al., 2018), and *SlHDT1* (Guo et al., 2022) were consistently documented to impede fruit ripening through the negative regulation of carotenoid accumulation. Additionally, SlHDA1 and SlHDA3 are engaged by other proteins during tomato fruit ripening, such as the interaction between SlERF.F12 and TOPLESS 2; moreover, the formation of a trimer involving SlHDA1/SlHDA3 inhibits fruit ripening (Deng et al., 2022).

We conducted in vitro experiments, such as Y2H (Fig. 5B) and Pull-down (Fig. 5C), to investigate whether the Thr mutation to Ser in *slwrky80^cr^*-46 leading to fruit browning is associated with SlHDAs and confirm the protein-level interaction between SlWRKY80 and SlHDA1. The predictions made by Alphafold2 (Fig. 5D) and Prodigy analysis (Supplementary table 3) indicate that after a single amino acid mutation in SlWRKY80, the protein structure interacting with SlHDA1 undergoes significant changes. As a result, its protein binding affinity and the dissociation constant at 25°C is increased, leading to a notable weakening of the interaction between the two proteins (Jumper et al., 2021). Concomitantly, the relative expression level of *SlHDA1* in the *slwrky80^cr^*-46 lines is notably lower than that in any other lines (Fig. 5E). The activity of SlHDA1 and the relative expression level of SlHDA1 have a similar trend (Supplementary figure 8), but the difference is that the *slwrky80^cr^*-46 line is significantly higher than the *slwrky80^cr^*-3 and *slwrky80^cr^*-4 lines. The D-Luc assay indicated that after the amino acid T-S mutation of SlWRKY80 (*slwrky80^cr^*-46 and *slwrky80^cr^*-T-S), its regulation of the *SlSGR1* promoter diminishes further in the presence of SlHDA1, elucidating further the significance of the weakened interaction intensity and its potential correlation with the color transition of tomato fruits. However, its regulation of the *SlPSY1* promoter significantly increases, leading to the suppression of chlorophyll degradation in tomato fruits and the promotion of phytoene synthesis. This phenomenon results in a brown coloration similar to that observed in the chlorophyll-deficiency *gf* mutant in tomato fruits (Fig. 5F). Thus, we hypothesize that as a regulatory factor, SlHDA1 is recruited by *SlWRKY80* to participate in fruit ripening and production of brown fruits. The study on the impact of changes in this type of amino acid on the accumulation and color transition of plant carotenoids was reported in *A. thaliana* (Yuan et al., 2015), *Armeniaca vulgaris* (Ni et al., 2022), and *S. lycopersicum* (Gady et al., 2012).

In summary, overexpressing of *SlWRKY80* leads to a reduction in the duration ofthe tomato color transition period. *S*lWRKY80 positively regulates the expression of downstream *SlSGR1* and *SlPSY1*. Promoting the breakdown of chlorophyll a and the synthesis of carotenoids, thereby expediting the overall color transition. SlHDA1 was recruited as a regulatory factor by SlWRKY80. The Thr mutation at the 69^th^ amino acid position of SlWRKY80 to Ser promoted the regulation of *SlPSY1*, inhibited the expression of *SlSGR1*, and resulted in tomato fruits exhibiting a brownish color similar to the green mutant gf.

## Materials and methods

### Plant materials

Wild type ‘Ailsa Craig’ (AC) tomato were used, while was used as the background material to construct *SlWRKY80* overexpressed lines (80OE-1 and 80OE-3) and *SlWRKY80* knockout lines (*slwrky80^cr^*-3, *slwrky80^cr^*-4, *slwrky80^cr^*-46). Overexpression vectors were constructed from tomato AC and pHellsgate2 (CaMV35S promoter driver) overexpressing *SlWRKY80* by cloning the CDs with SmaI and KpnI (modified by Prof. Zhan Xiangqiang, College of Horticulture, Northwest A&F University). Predicting SlWRKY80 target information through CRISPR RGEN Tools (http://www.rgenome.net/casdesigner/result?hash=49d4ef08e39c96a14781bc8f463be7f6) and constructing CRISPR/Cas9:*SlWRKY80*. Sequencing was performed by Sangon Biotech (Shanghai, China). The *SlWRKY80* from different genotypes was compared with DNAMAN v.6 (Lynnon Biosoft, CA, USA). The primers used are listed in Supplementary table S4.

The aforementioned recombinant vectors were utilized to obtain transgenic materials via Agrobacterium-mediated infection (Xu et al., 2023). Identification was performed using specific primers (Supplemental table S4), and positive materials were retained for seed collection and continued propagation. The T2 generations of homozygous overexpressed (80OE-1, 80OE-3) and knockout lines (*slwrky80cr*-3, *slwrky80cr*-4, *slwrky80cr*-46) were obtained by self-crossing to obtained transgenic lines.

### Plant growth conditions

Seedlings were cultivated in a growth chamber (GXZ-5000E, China Ningbo Southeast Instrument Co., Ltd.) with a 200 μmol·m^−2^·s^-1^, a 12-hour daily light period, temperature and humidity are 25 ℃ and 20 ℃ (day and night).

### RNA extraction and Real-Time quantitative PCR (RT-qPCR)

Select the sampling location for the fruit pulp, as shown in the red circle in Figure 3A for specific locations. RNA extraction and first-strand cDNA synthesis and RT-qPCR were carried out following the protocol established by Xu et al (2023) and Livak et al (2001). Related primers provided in Supplementary table S5.

### Determination of chlorophyll and carotenoids, etc

Chlorophyll a content, chlorophyll b content, and carotenoids are determined by extraction method (Costache et al., 2012). The responsibility of determining the LC-MS/MS method for β-carotenoids, lutein, and lycopene was conferred upon Wuhan Maiteweier Biotechnology Co., Ltd. (Wuhan, China) (Petry et al. 2018). The above data is consistent with the sampling site and RNA sampling site.

### Determination of fruit hardness, lycopene content and soluble solids content

Fruit firmness was determined using a digital force pressure tester equipped (Model Instron 5542, Instron Co., USA). The content of lycopene was determined using the method proposed by Davis et al. (2003), while the content of soluble solids was determined using the method proposed by Yang et al. (2009).

### SlHDA1 activity determination

Take 0.1g of tomato pulp, freeze it in liquid nitrogen and grind it into powder. Add 500 μl of diluted 1 × NE1 (containing dithiothreitol/DTT), thoroughly vortex and mix well. Follow the instructions in the Plant histone deacetylase 1 (HDAC1) ELISA scientific research kit (MM-6378302) for the subsequent steps. The extracted nuclear protein was measured for initial protein concentration using an Infinite M200pro and adjusted for consistency with ddH2O. The protein was stored at −80℃ for later use. The SlHDA1 activity detection kit was purchased from Jiangsu Meimian Industrial Co., Ltd.

### Transmission electron microscopy (TEM)

The TEM experiment was commissioned by Wuhan Saiweier Biotechnology Co., Ltd. (Wuhan, China), the organizational position used is the flesh part of tomato fruit (shown in the red circle in Figure 3A).

### Yeast one-hybrid (Y1H) assays

*SlWRKY80* cDNA was sequenced and cloned into pGADT7 vector for construction of prey-*SlWRKY80*. Insert the 6 promoter sequences of *SlSGR1* and the 2 promoter sequences of *SlPSY1* into the pAbAi vector respectively to construct the pBait reporter vector. Upon completing vector construction, the Y1H procedure was conducted in accordance with the methodology outlined by Liang et al. (2023). Supplementary table S4 lists the related primers.

### Electrophoretic mobility shift assay (EMSA)

The *SlWRKY80*’s CDs was cloned into the pGEX4T-1 expression vector using the *EcoR* I and *Sal* I sites to produce a glutathione S-transferase fusion protein. Related primers can be found in supplemental table S4. The fusion protein was then transformed into *Escherichia coli* lines BL21, and induction was carried out using constant shaking at 200 r/min and a temperature of 28°C for 8 hours with an IPTG concentration of 0.5 mM. A glutathione sepharose beads (635608, Takara)-based purification method was then used to purify the fusion protein. Invitrogen synthesized the biotin-labeled *SlSGR1* and *SlPSY1* promoter oligonucleotide probes, which are listed in supplemental table S4. EMSAs were performed utilizing a LightShift Chemiluminescent EMSA Kit (Beyotime). The mutation and cold probes employed for EMSA can also be found in supplemental table S6.

### Yeast two-hybrid (Y2H) assay

To assess the self-activation of the *SlWRKY80*, we divided it into two fragments consisting of a conserved domain, which were subsequently cloned separately into the yeast pGBKT7 vector, with BD-*SlWRKY80*_68-273_, and BD-*SlWRKY80*_1-273_ cleavage sites. Supplemental table S4 lists the primer sequences. The BD-*SlWRKY80*_68-273_, BD-*SlWRKY80*_1-273_ and pGADT7 were transformed into Y2H. After transformation, 5 µL of the transformed cells were placed onto solid-state media of two deficient DDO (SD/- Trp/-Leu) and four deficient QDO (SD/- Trp/Leu/- Hiss/- Ade). 3-5 days were spent incubating the media at 28℃, and yeast growth was recorded. From this, it can be concluded that BD-*SlWRKY80*_68-273_ is a non self activating fragment containing the structural domain of *SlWRKY80*.

The CDs of 13 genes from the SlHDA family in tomato were cloned into yeast pGADT7 vector with *Nde*I and *Xho*I cleavage sites, and then co transformed with BD-*SlWRKY80*_68-273_ into Y2H. After transformation, 5 µL of the transformed cells were placed onto solid state media of two defective DDO and four defective QDO. The media was incubated at 28℃ in an incubator for 3-5 days.

### Pull-down

The induction and purification of the GST-*SlWRKY80* fusion protein can be found in the EMSA test. Cloning of the coding sequence (CDS) of *SlHDA1* into pMAL-C2x was performed using *EcoR*I and *Hind*III. The expression vector was generated using relevant primers (Supplementary table S4) to produce the MBP-*SlHDA1* fusion protein. Next, the *E. coli* BL21 lines were transformed with the fusion protein and grown by shaking the culture at 200 r/min and 28°C for 8 hours, with induction induced using IPTG at a concentration of 0.5 mM. The subsequent removal of the medium was performed by centrifugation (A temperature of 4°C, a speed of 4000 rpm, and a duration of 10 minutes), and the supernatant precipitate was discarded. The remaining substance was resuspended in 1xPBS (pH=8.0) and subsequently subjected to ultrasound treant on ice for 15 minutes. The solution was then centrifuged again (A temperature of 4°C, a speed of 4000 rpm, and a duration of 10 minutes), and collecting the supernatant. The maltose binding protein label protein purification kit was used to further purify the obtained MBP-*SlHDA1* fusion protein (Abbkine (Wuhan) Biotechnology Co., Ltd., Wuhan, China).

The GST Pull-down analysis experience refers to the method of Zhang et al. (2023).

### Dual-luciferase (LUC) analysis

*SlWRKY80*’s full-length CDs, excluding the termination codon, was cloned into the pGreenⅡ-002962-SK vector using *Sac*I and *Kpn*I cleavage sites. The pro-SlSGR1-P2/6 and pro-SlPSY1-P1/2 were constructed on pGreen II-0800 vector through homologous recombination to determine the strength of each promoter of SlSGR1 and SlPSY1 binding to SlWRKY80 protein. The upstream 3000bp of SlSGR1 and SlPSY1 promoters were constructed on pGreen II-0800 vector through homologous recombination (Report1 and Report2 in Fig. 5F) to investigate the effect of SlHDA1 on the regulation of downstream SlSGR1 and SlPSY1 genes by SlWRKY80. In supplemental table S4 are listed the primers used. The D-LUC test operation is based on the method of Liang et al. (2023). The tobacco plants were then incubated for 72 hours before fluorescence detection was performed using the Dual Lucifera Reporter Assay System (E1910, Promrga, USA).

### Statistical analysis

LSD’s test, based on DPS7.5, was used to detect significant differences between three replicates of each experiment.

## Acknowledgements

This work supported by China Agriculture Research System (CARS-23-D06). In addition, we thank Professor Jiafa Wang for providing pRSFDuet-1 engineering bacteria, we thank Professor Zhan Xiangqiang for providing the pHellsgate2 carrier.

## Author contributions

X.H. and C.S. conceived and designed the experiments. C.S., G.C., and X.L. wrote the paper. C.S., G.C., Z.L., and H.Z. performed the experiments. G.L., S.H., J.W., and X.H. revised the manuscript and edited the final version. All authors read and approved the manuscript for submission.

## Supplemental data

The following materials are available in the online version of this article.

**Supplement figure 1**. Analysis of W-box cis acting elements in *SlSGR1* and *SlPSY1* promoters.

**Supplementary figure 2.** The content of carotenoid metabolism pathway related substances in tomato fruits of WT, *slwrky80cr*-4 and *slwrky80cr*-46 lines during the Break and Break+7 stages.

**Supplementary figure 3.** GO classification diagram of the comparison between the fruits of WT, *slwrky80cr*-4 and *slwrky80cr*-46 lines during Break+7 period.

**Supplement table 1.** Days from anthesis to breaker stage for Wild type and SlWRKY80 transgenic lines.

**Supplement table 2.** Analysis of phosphorylation sites near the 69th position of different amino acid editing types in SlWRKY80

**Supplement table 3.** Alphafold2 predicts protein-protein interaction related indicators.

**Supplement table 4.** Rt-PCR Primers

**Supplement table 5.** Rt-qPCR Primers

**Supplement table 6.** The mutation and cold probes

## Data availability

The authors affirm that all the information required to substantiate the findings of the study is provided in both the paper and the supplementary materials, or can be acquired by contacting the corresponding author.

## Antibodies resources

Anti-Flag was purchased from Sangon Biotech; anti-GST and anti-MBP antibodies were purchased from Abcam.

## Accession numbers

A database of online sequence data was used to obtain sequence data (https://solgenomics.net/search/locus?tdsourcetag=spcqq_aiomsg).

The gene accession numbers used in this study can be found below : Solyc03g095770.2 (*SlWRKY80*), Solyc08g080090.2 (*SlSGR1*), Solyc03g031860.2 (*SlPSY1*), Solyc11g066440.1 (*SlPAO*), Solyc01g088090.2 (*SlPPH*), Solyc07g024000.2 (*SlNYC1*), Solyc09g091440.3 (*SlHDA1*), Solyc03g112410.2 (*SlHDA2*), Solyc06g071680.3 (*SlHDA3*), Solyc11g067020.2 (*SlHDA4*), Solyc08g065350.3 (*SlHDA5*), Solyc06g074080.3 (*SlHDA6*),S olyc01g009110.2 (*SlHDA7*), Solyc03g119730.3 (*SlHDA8*), Solyc03g115150.3 (*SlHDA9*), Solyc01g009120.3 (*SlHDA10*), Solyc09g009030.3 (*SlHDT1*), Solyc10g085560.2 (*SlHDT2*), Solyc11g066840.2 (*SlHDT3*).

---

**Supplementary figure 1.**
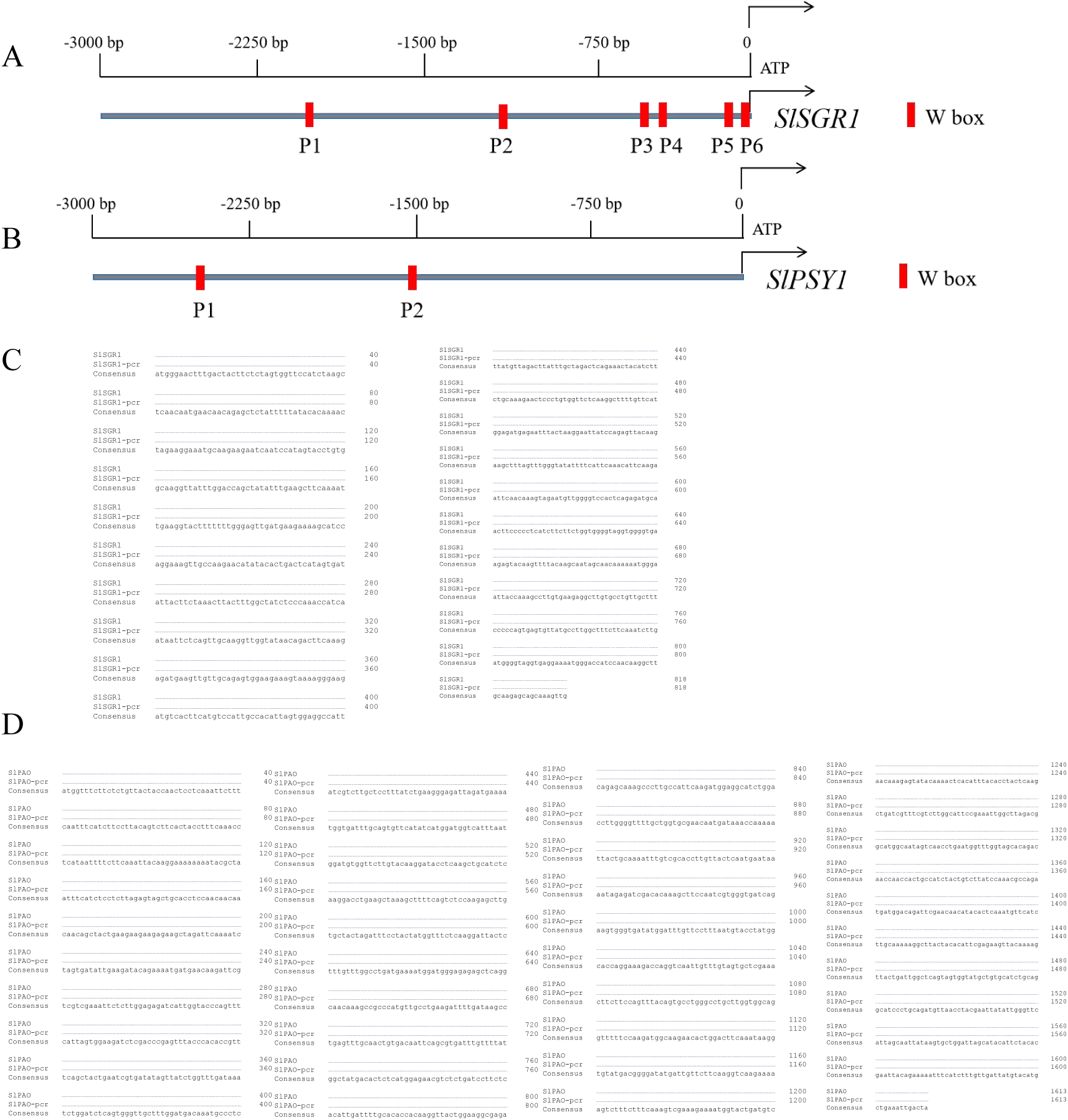
Analysis of W-box cis acting elements in the promoters of *SlSGR1* and *SlPSY1*, as well as sequencing comparison of *SlSGR1* and *SlPSY1* in the WT and *slwrky80^cr^*-46 strains. A and B, W-box analysis of upstream 3000bp sequences of *SlSGR1* and *SlPSY1* promoters. C and D, Sequencing comparison of *SlSGR1* and *SlPSY1* in WT and slwrky80cr-46 strains.

**Supplementary figure 2.**
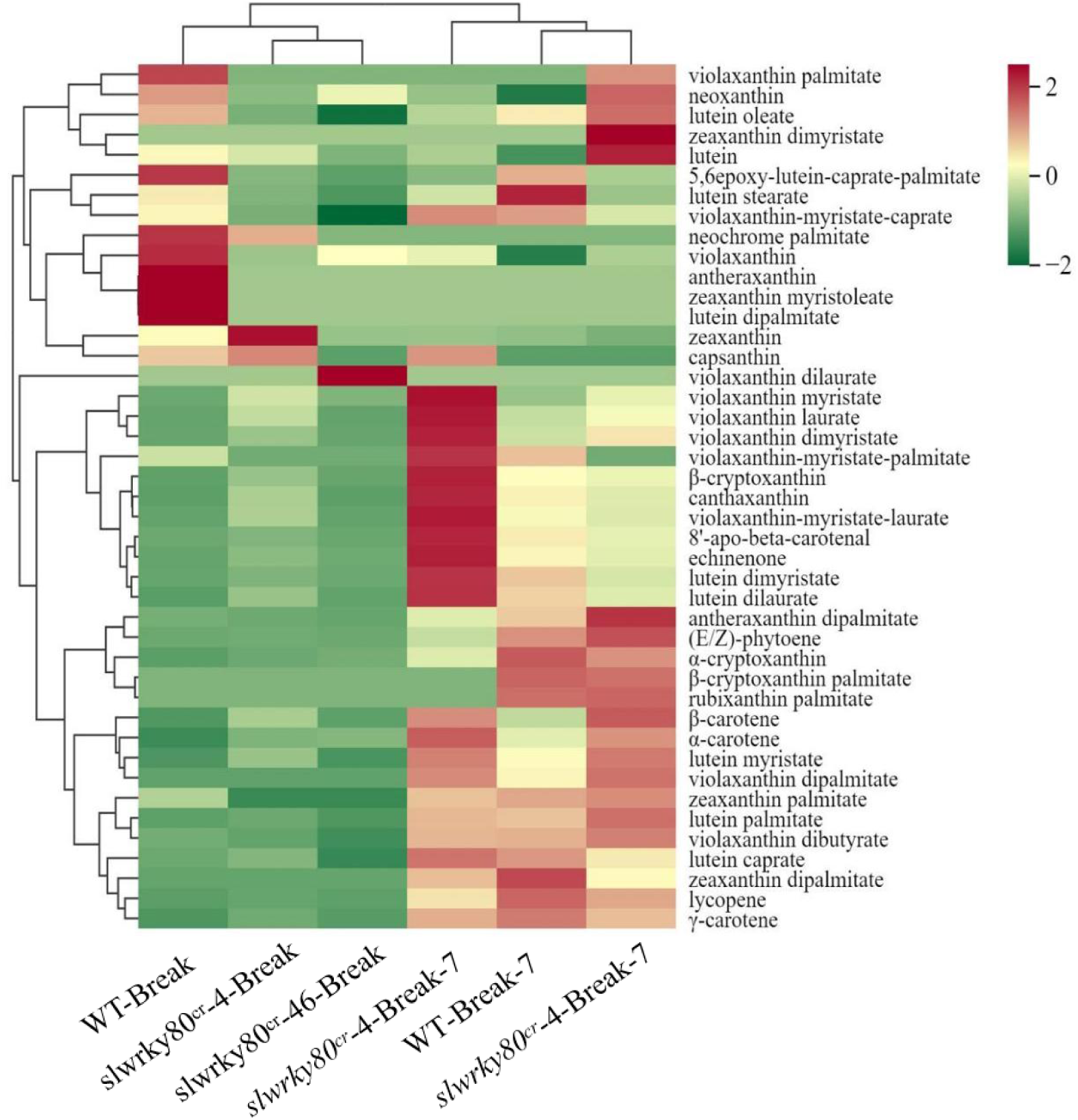
The content of carotenoid metabolism pathway related substances in tomato fruits of WT, *slwrky80^cr^*-4 and *slwrky80^cr^*-46 lines during the Break and Break+7 stages.

**Supplementary figure 3.**
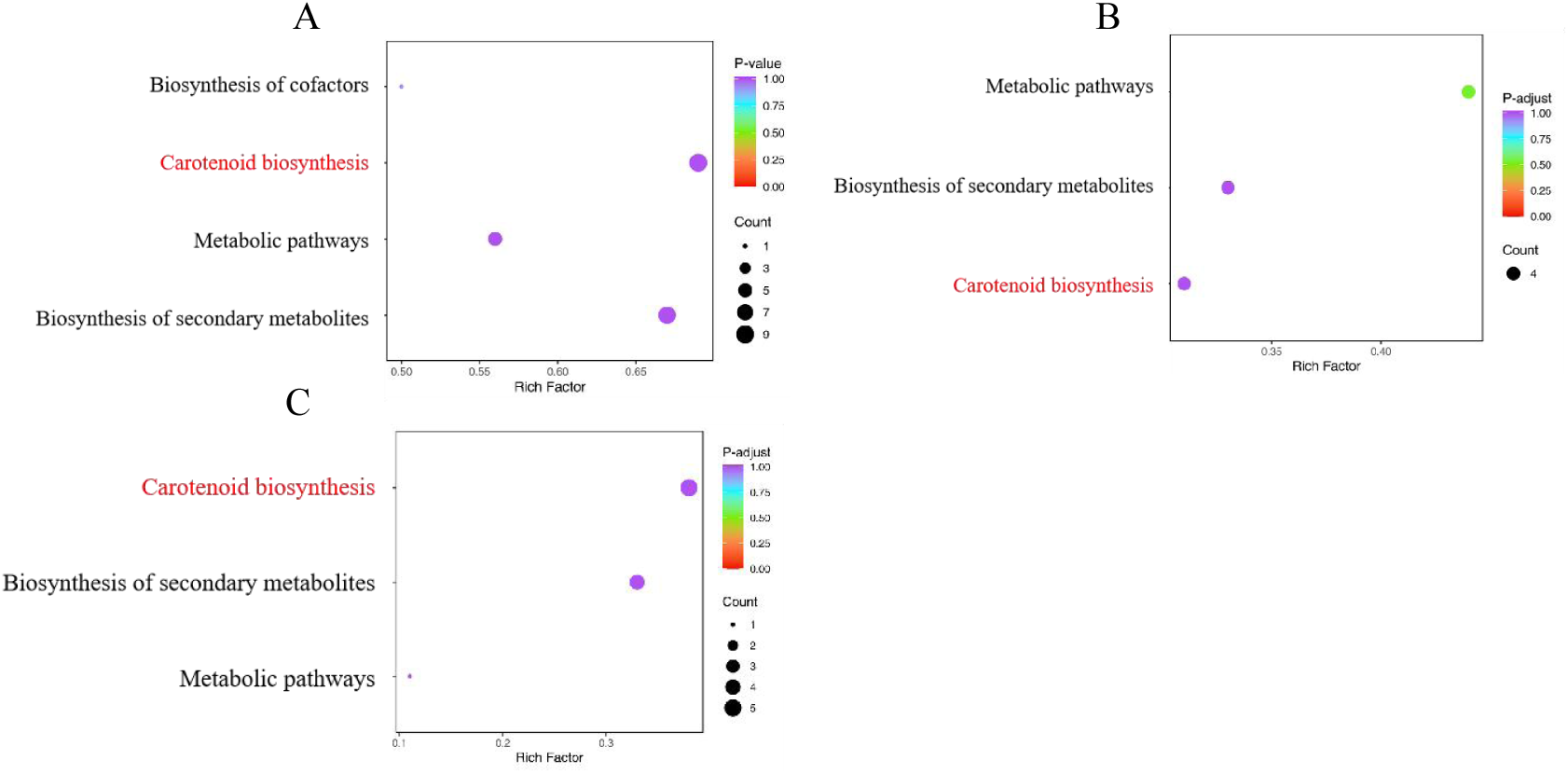
GO classification diagram of the comparison between the fruits of WT, *slwrky80^cr^*-4 and *slwrky80^cr^*-46 lines during Break+7 period. A, *slwrky80^cr^*-4-Break+7 vs WT-Break+7; B, *slwrky80^cr^*-46-Break+7 vs WT-Break+7; C, *slwrky80^cr^*-46-Break+7 vs *slwrky80^cr^*-4-Break+7.

**Supplementary figure 4.**
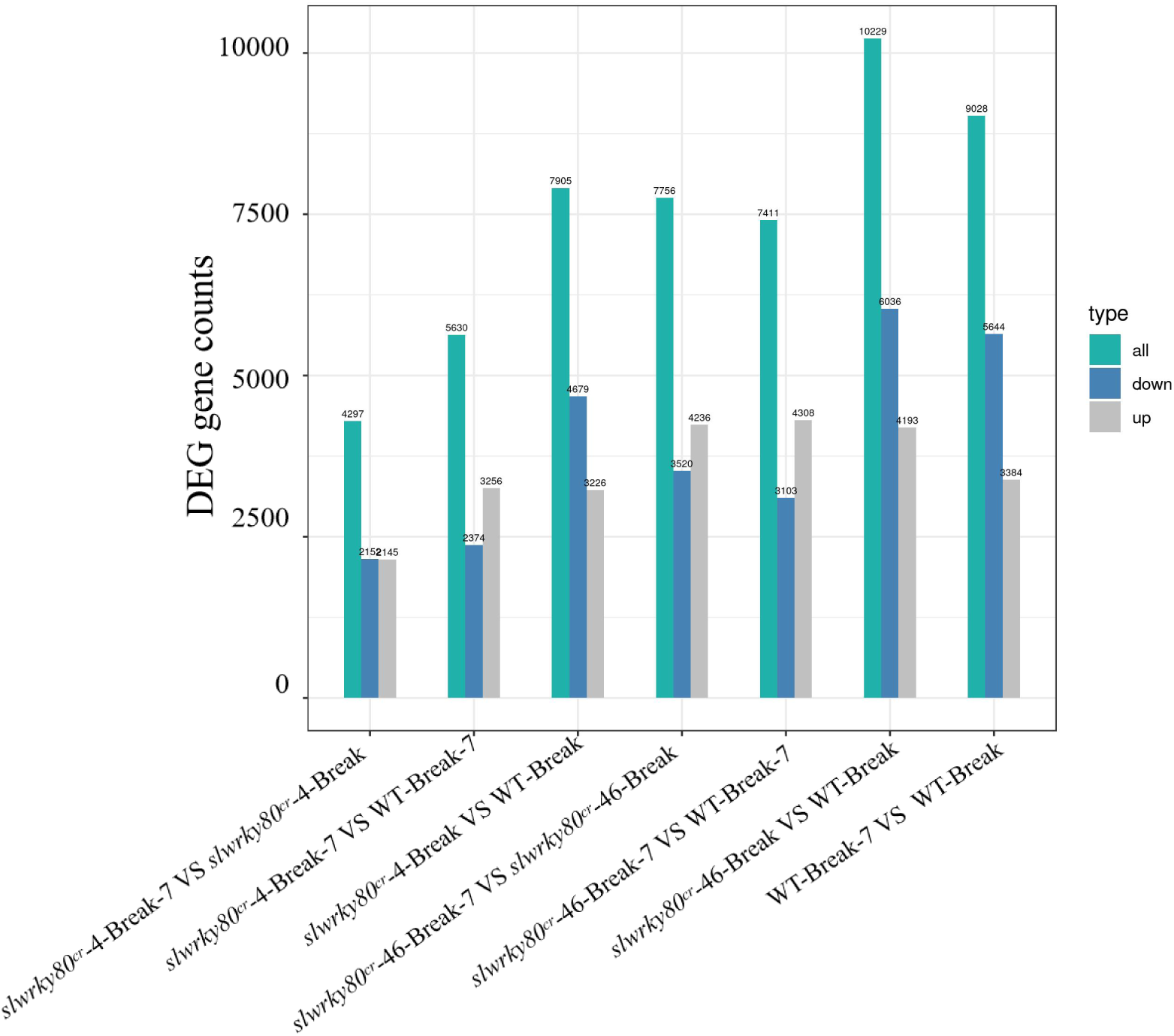
The DEG gene counts in tomato fruits of WT, *slwrky80^cr^*-4 and *slwrky80^cr^*-46 lines during the Break and Break+7 stages.

**Supplementary figure 5.**
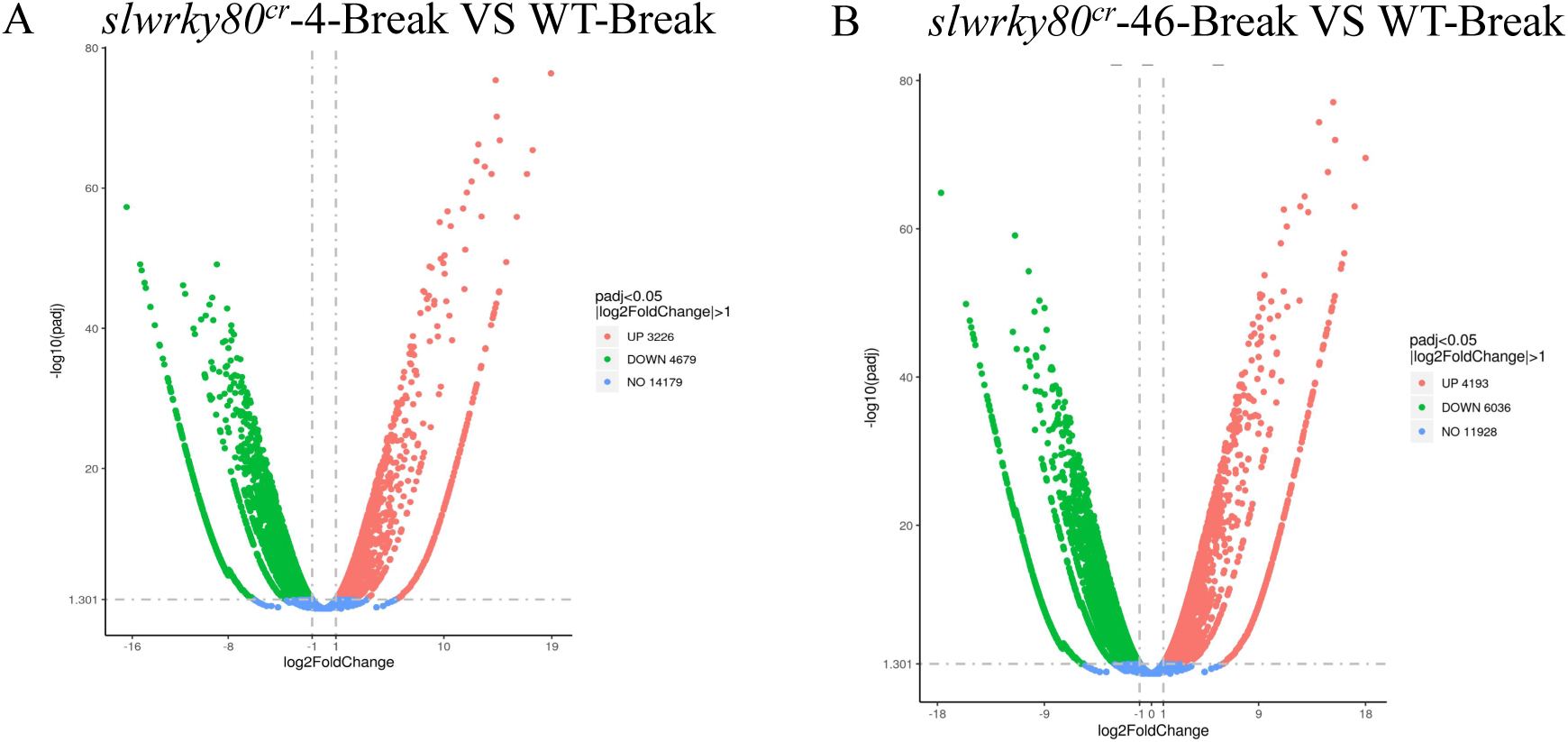
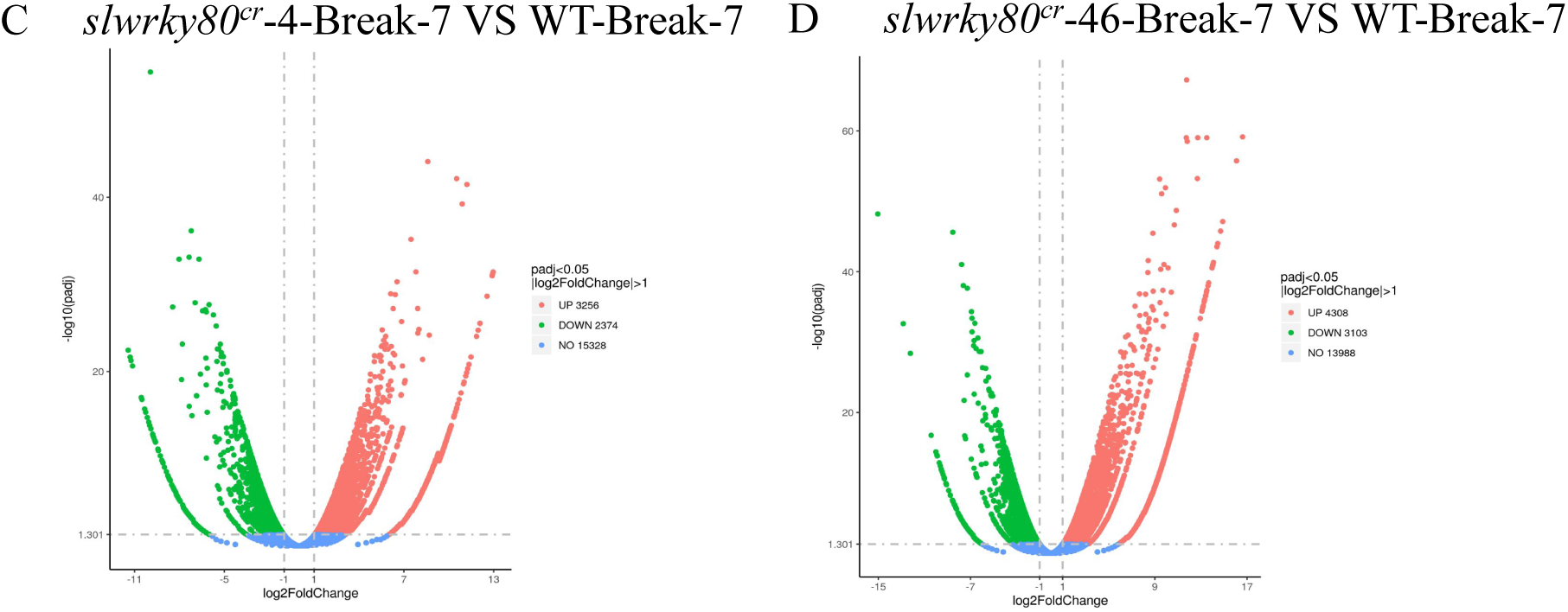
Gene volcano maps of *slwrky80^cr^*-4 and *slwrky80^cr^*-46 lines compared to WT during Break and Break+7 periods. A, *slwrky80^cr^*-4-Break VS WT-Break. B, *slwrky80^cr^*-46-Break VS WT-Break. C, *slwrky80^cr^*-4-Break-7 VS WT-Break-7. D, *slwrky80^cr^*-46-Break-7 VS WT-Break-7.

**Supplementary figure 6.**
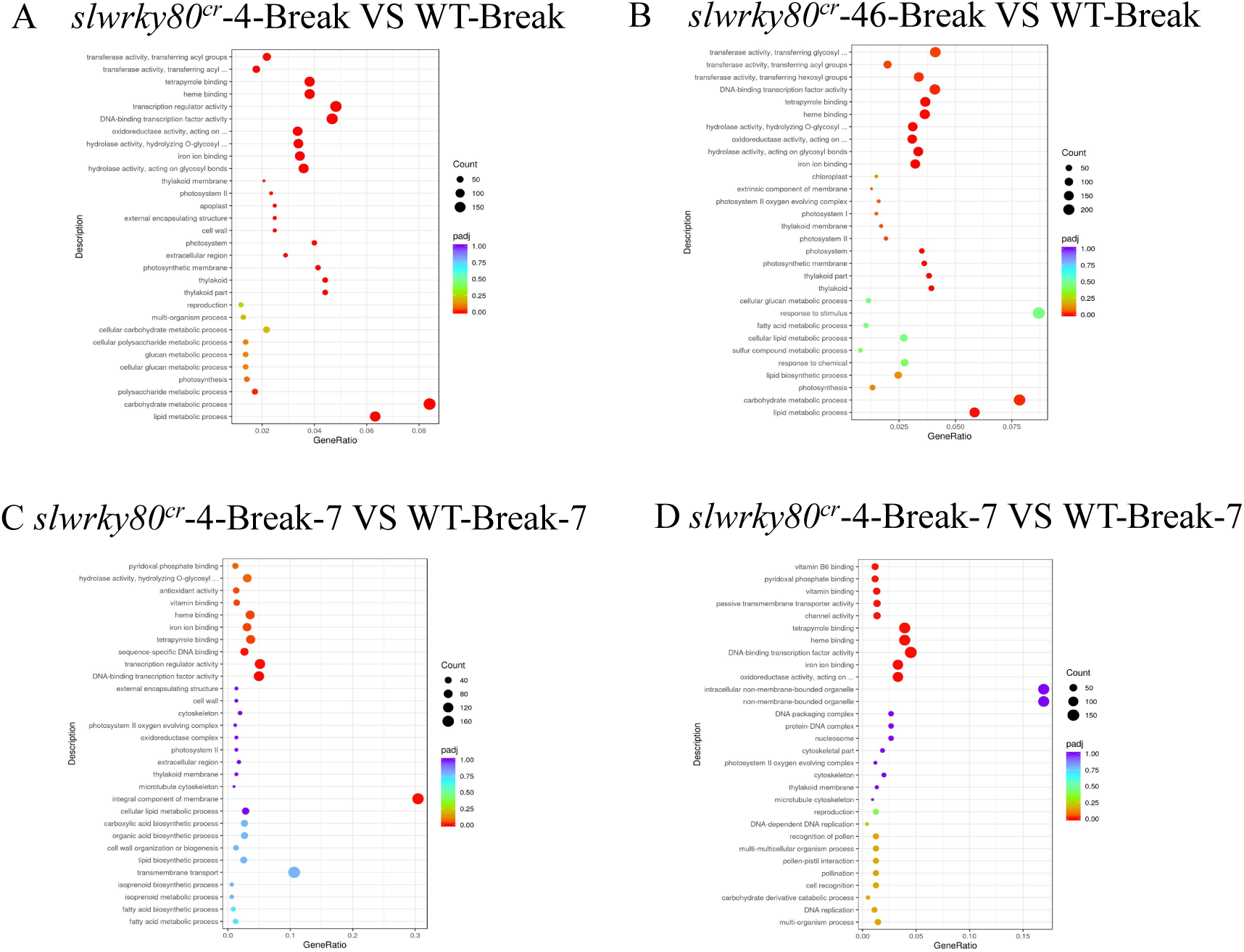
GO classification map of *slwrky80^cr^*-4 and *slwrky80^cr^*-46 lines compared to WT during Break and Break+7 periods. A, *slwrky80^cr^*-4-Break VS WT-Break. B, *slwrky80^cr^*-46-Break VS WT-Break. C, *slwrky80^cr^*-4-Break-7 VS WT-Break-7. D, *slwrky80^cr^*-46-Break-7 VS WT-Break-7.

**Supplementary figure 7.**
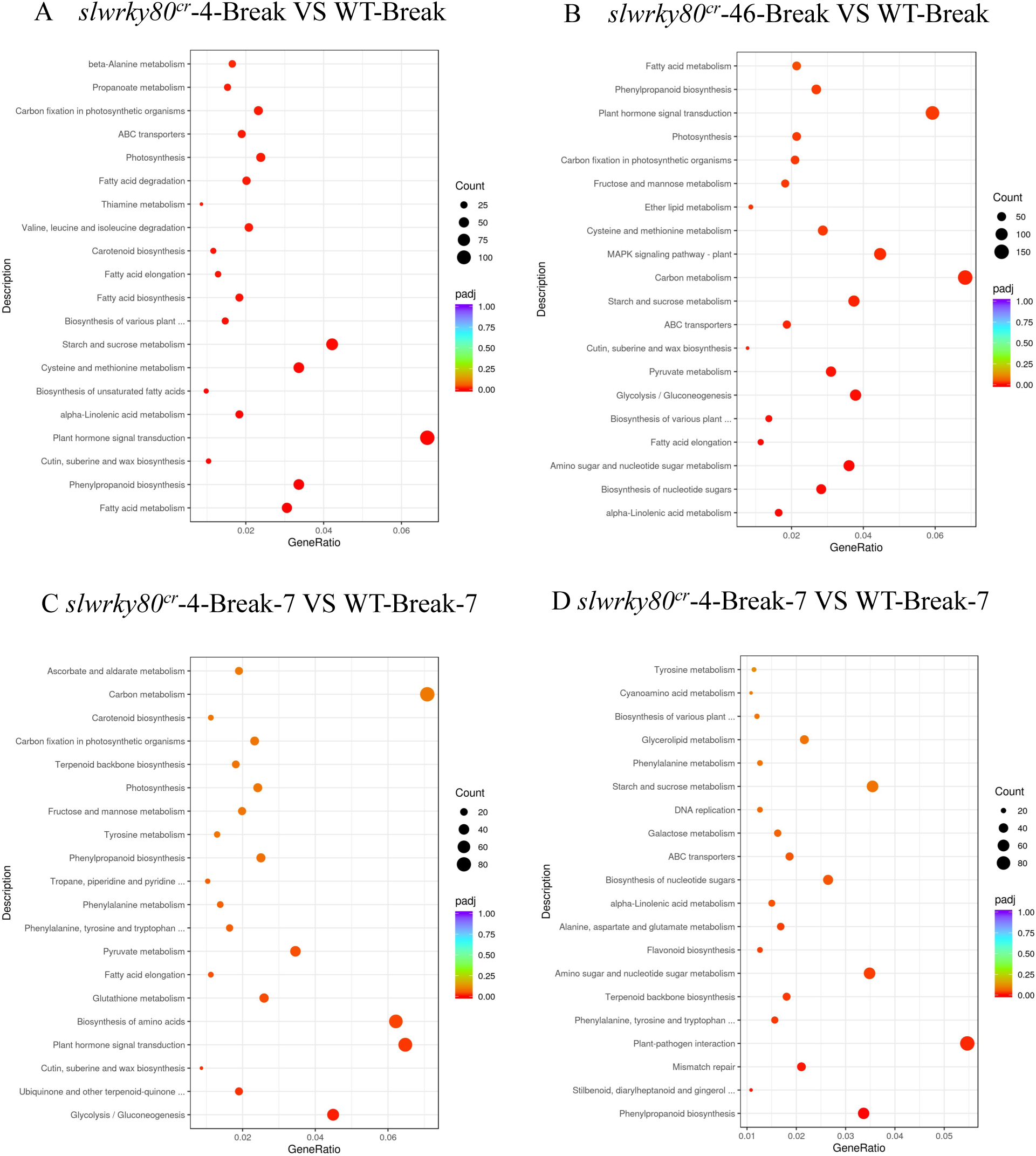
KEGG map of *slwrky80^cr^*-4 and *slwrky80^cr^*-46 lines compared to WT during Break and Break+7 periods. A, *slwrky80^cr^*-4-Break VS WT-Break. B, *slwrky80^cr^*-46-Break VS WT-Break. C, *slwrky80^cr^*-4-Break-7 VS WT-Break-7. D, *slwrky80^cr^*-46-Break-7 VS WT-Break-7.

**Supplementary figure 8.**
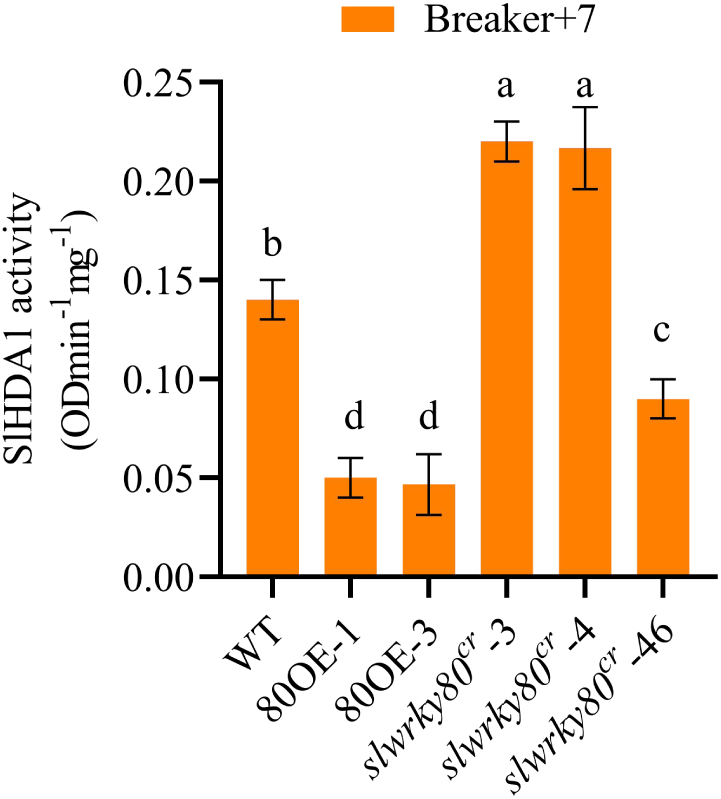
SlHDA1 activity at Break+7 period. Values represent the mean across three biological replicates, error bars represent SD, and significant differences (*P*<0.05) are indicated by lowercase letters based on LSD’s test.

**Supplement table 1.**
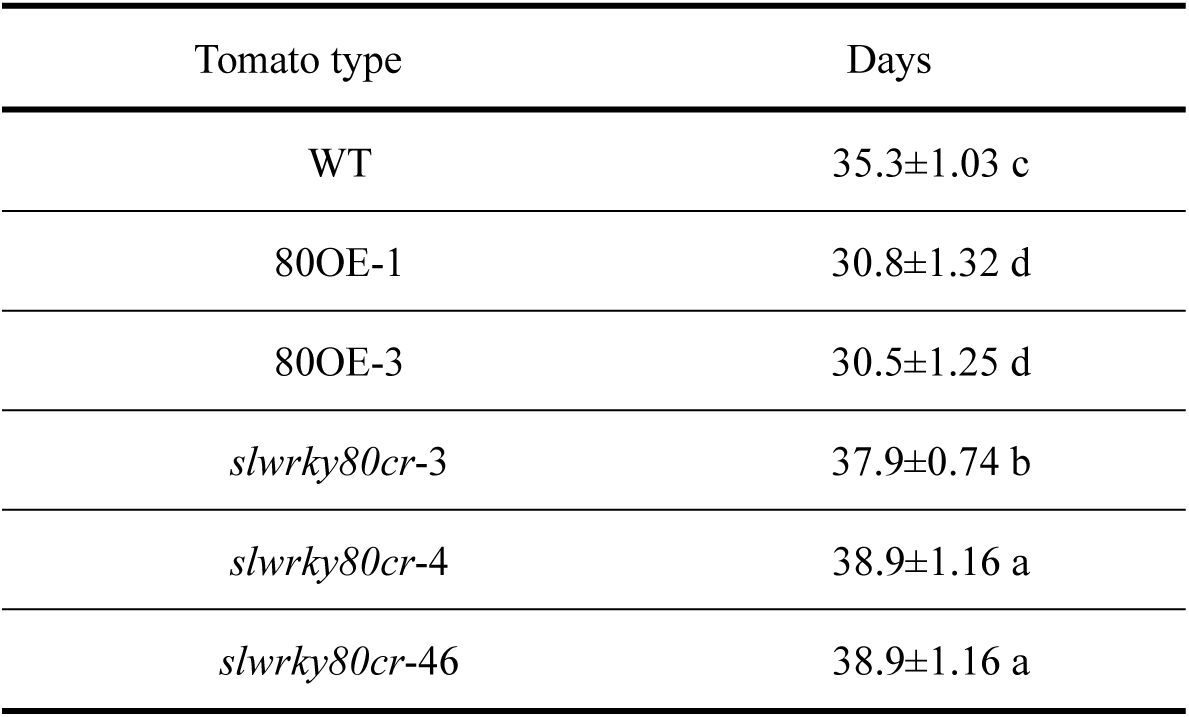
Days from anthesis to breaker stage for Wild type and SlWRKY80 transgenic lines. 15 fruits were examined for biological replicates per lines in Table 1.

**Supplement table 2.** Analysis of phosphorylation sites near the 69^th^ position of different amino acid editing types in SlWRKY80.

|  | No. | Amino<br>acid type | Context | Score | Kinase | Answer |
| --- | --- | --- | --- | --- | --- | --- |
| SIWRKY80 | 69 | T | DNSLT<br>QGQI | 0.520 | ATM | YES |
|  | 81 | S | DQPNS<br>GSEL | 0.982 | unsp | YES |
| *slwrky80cr-3* | 69 | T | DNSLT<br>QVEE | 0.782 | unsp | YES |
|  | 81 | S | PNSGSE<br>LKK | 0.614 | PKC | YES |
| *slwrky80cr-4* | 69 | T | DNSLTR<br>GSS | 0.475 | GSK3 | No |
|  | 72 | S | LTRGSS<br>TQF | 0.955 | unsp | YES |
| *slwrky80cr-46* | 69 | S | DNSLS<br>RKLI | 0.446 | GSK3 | No |
|  | 87 | S | RRRRS<br>KTVE | 0.998 | unsp | YES |
| SIWRKY80-T-S | 69 | S | DNSLS<br>QGQI | 0.843 | unsp | YES |
|  | 81 | S | DQPNS<br>GSEL | 0.982 | unsp | YES |

**Supplement table 3.** Alphafold2 predicts protein-protein interaction related indicators.

|  | Predicted binding affinity<br>(kcal.mol<sup>-1</sup>) | Predicted dissociation constant<br>(M) at 25.0°C |
| --- | --- | --- |
| SIWRKY80+SIHDA1 | -18.9 | 1.3e-14 |
| SIWRKY80-T-S+SIHDA1 | -14.5 | 2.2e-11 |

**Supplement table 4.**
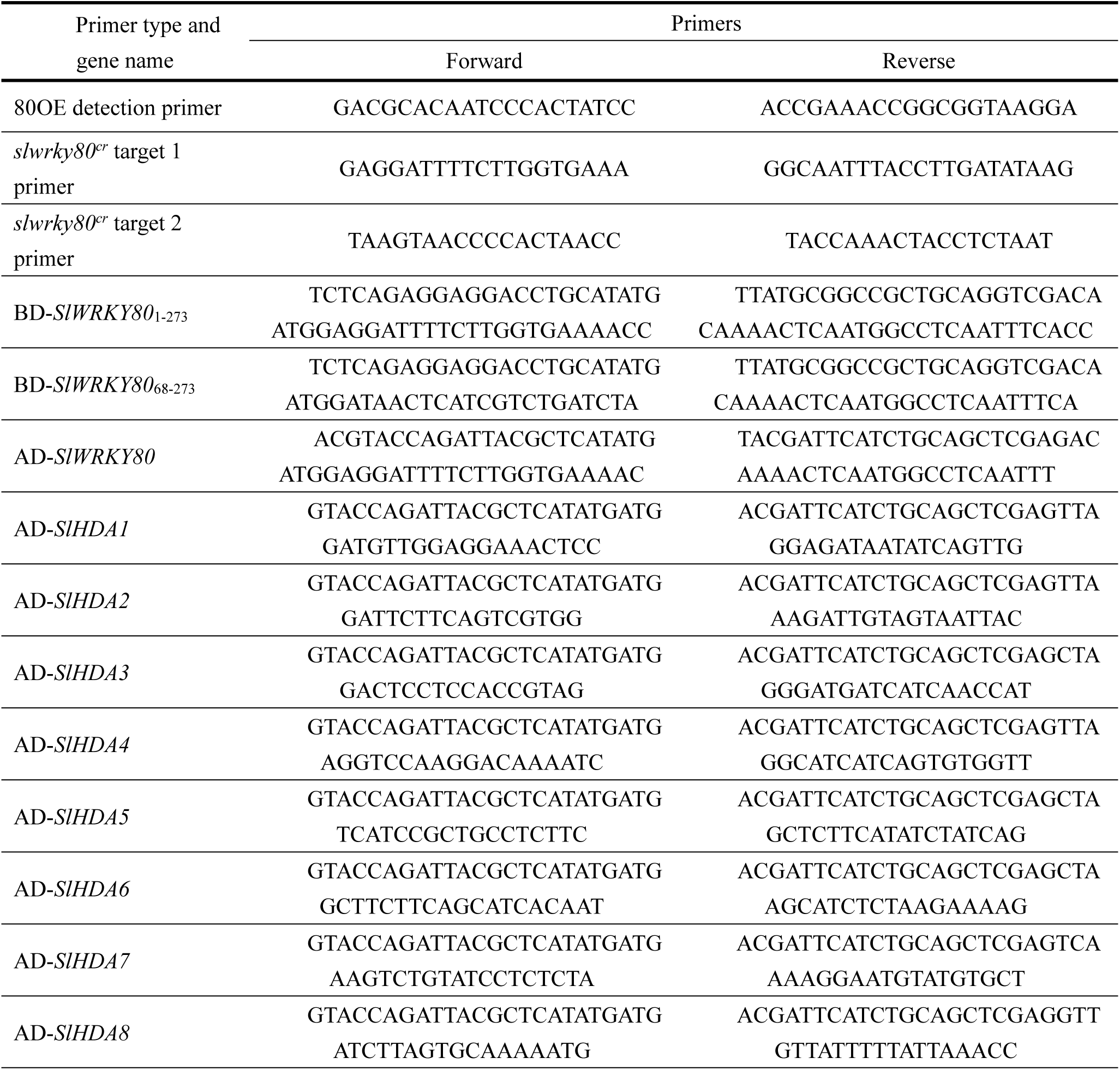
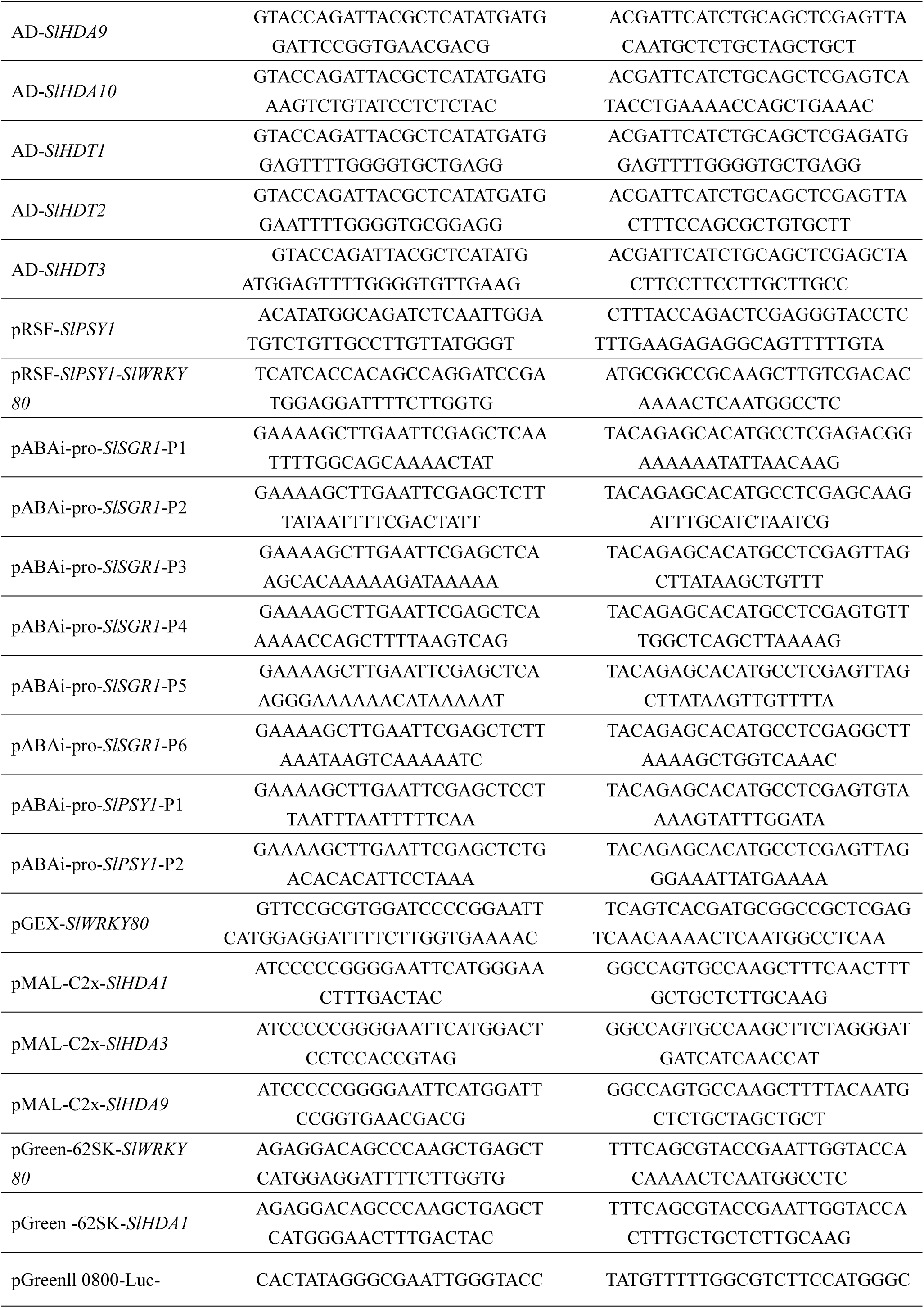
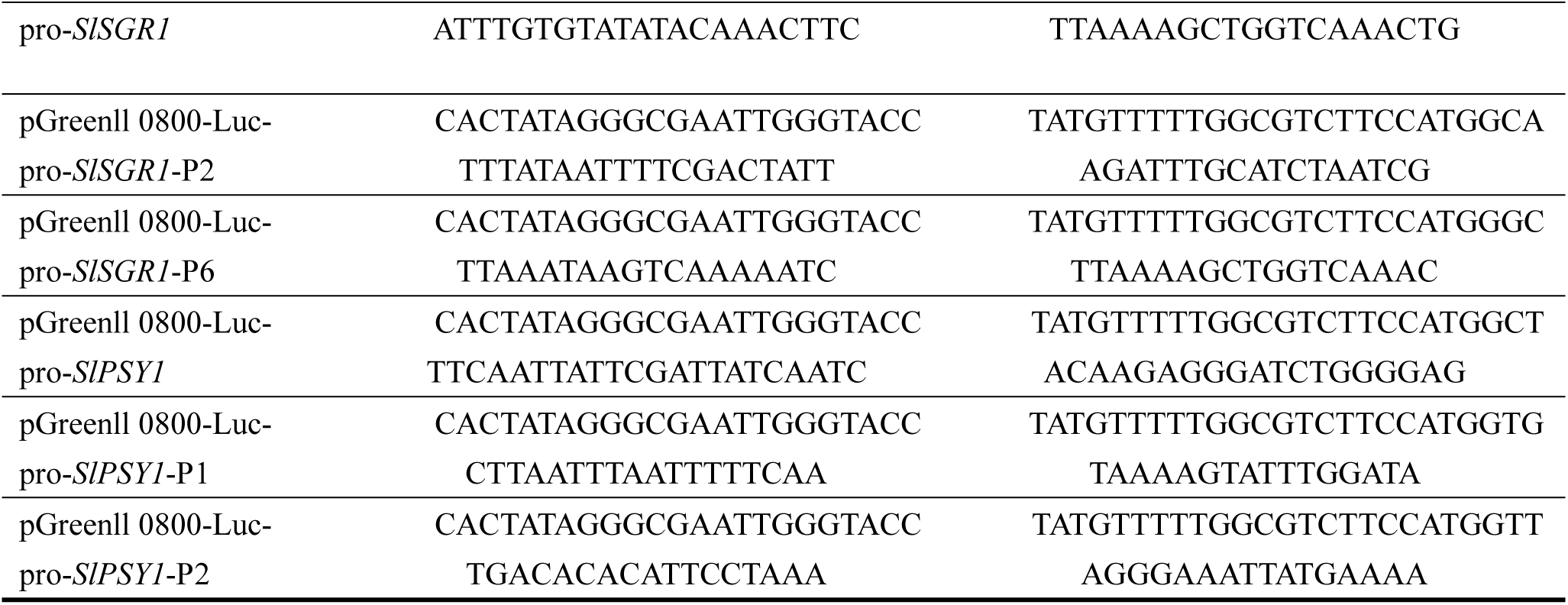
Rt-PCR Primers.

**Supplement table 5.**
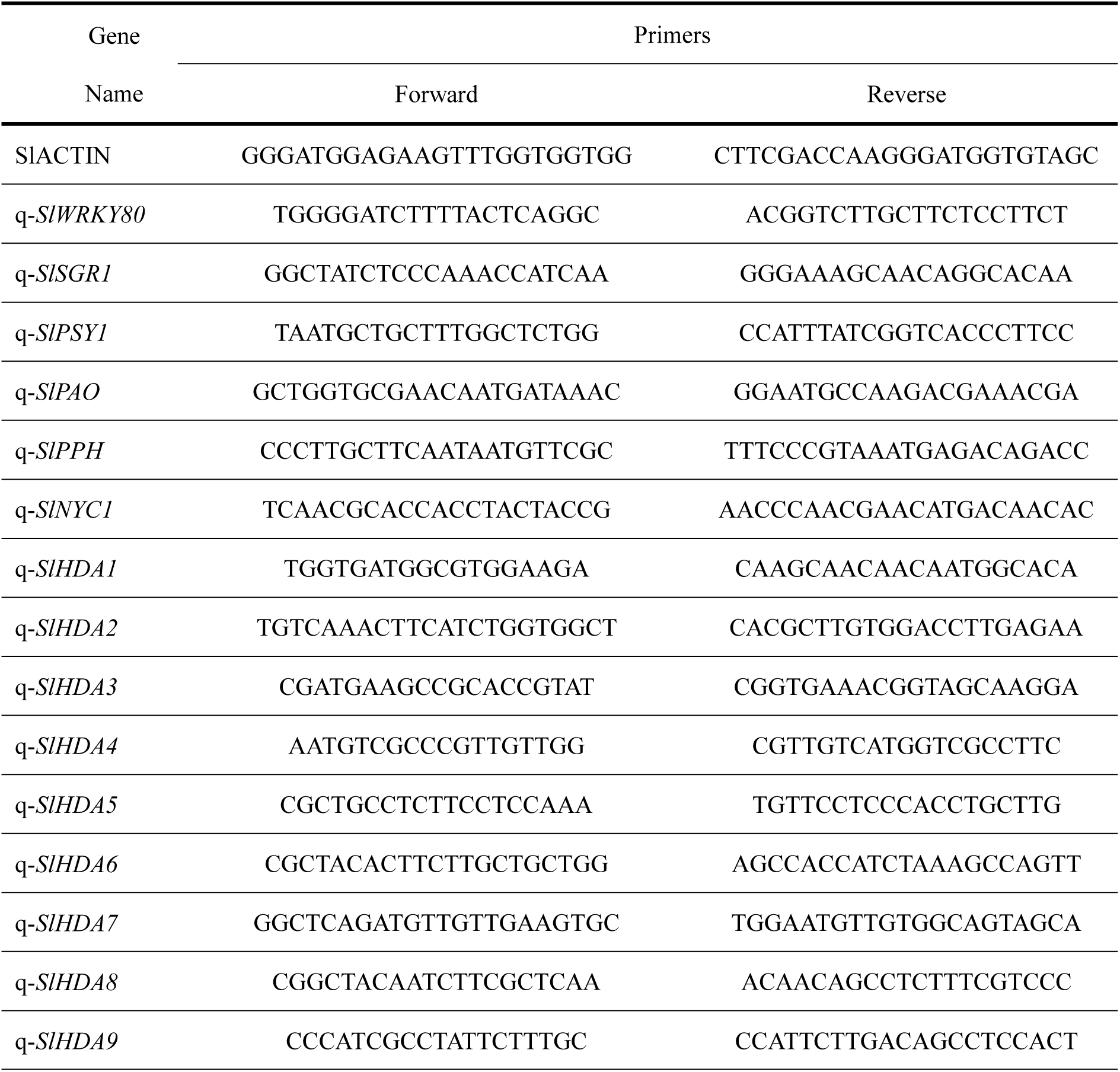
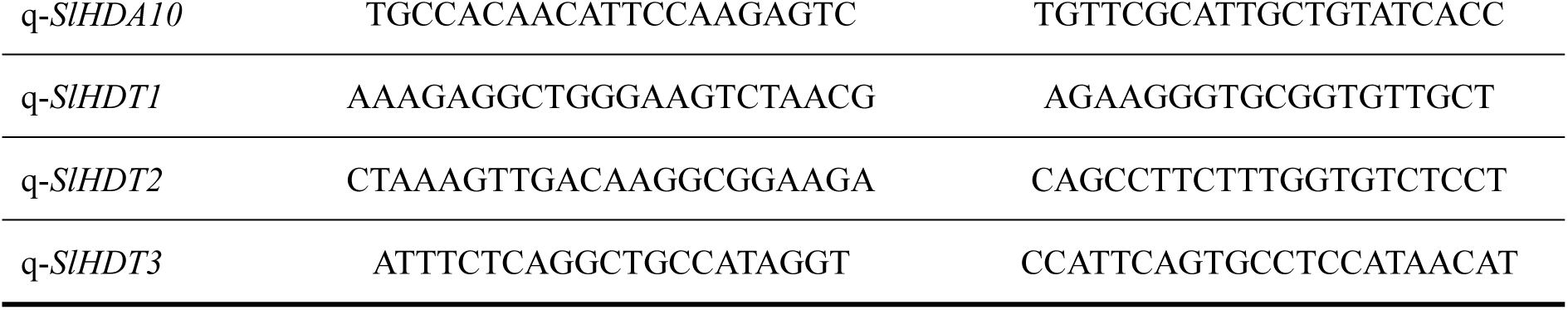
Rt-qPCR Primers.

**Supplement table 6.**
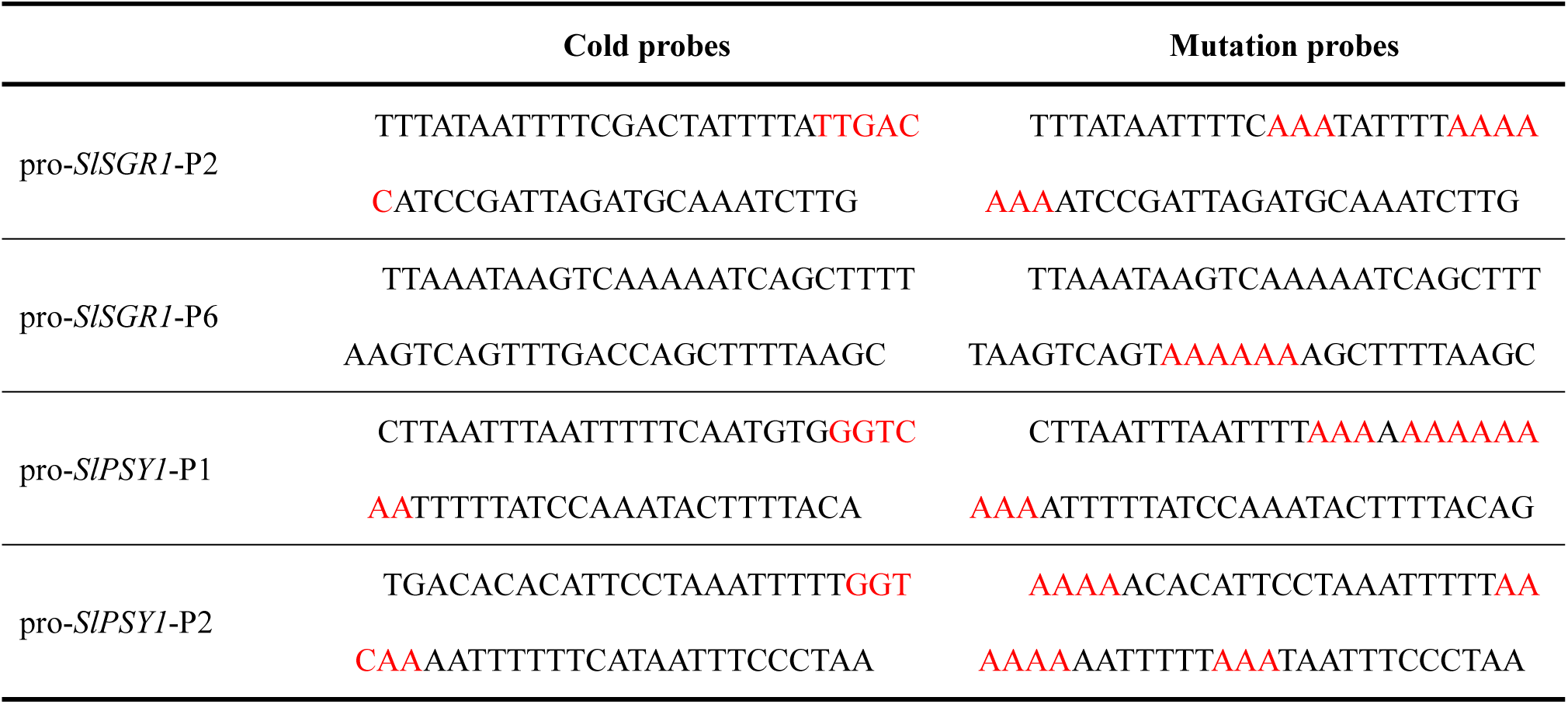
The mutation and cold probes.

## References

Akhtar, M, S., Goldschmidt, E, E., John, I., Rodoni, S., Matile, P., and Grierson, D. (1999). Altered patterns of senescence and ripening in gf, a stay-green mutant of tomato (*Lycopersicon esculentum* Mill.). J Exp Bot 50: 1115–1122.

Arhondakis, S., Bita, C, E., Perrakis, A., Manioudaki, M. E., Krokida, A., Kaloudas, D., and Kalaitzis, P. (2016) In silico transcriptional regulatory networks involved in tomato fruit ripening. Front Plant Sci 7: 1234.

Barry, C, S., Mcquinn, R, P., Chung, M, Y., Besuden, A., and Giovannoni, J, J. (2008). Amino acid substitutions in homologs of the STAY-GREEN protein are responsible for the green-flesh and chlorophyll retainer mutations of tomato and pepper. Plant Physiol 147: 179.

Costache, M, A., Campeanu, G., and Neata, G. (2012). Studies concerning the extraction of chlorophyll and total carotenoids from vegetables. Rom Biotech Lett 17: 7702–7708.

Davis, A, R., Fish, W, W., and Perkins-Veazie, P. (2003). A rapid spectrophotometric method for analyzing lycopene content in tomato and tomato products. Postharvest Biol Tec 28: 425–430.

Deng, H., Chen, Y., Liu, Z., Liu, Z, Q., Shu, P., Wang, R, C., Hao, Y, W., Su, D., Pirrello, J., and Liu, Y, S. (2022). SlERF.F12 modulates the transition to ripening in tomato fruit by recruiting the co-repressor TOPLESS and histone deacetylases to repress key ripening genes. The Plant Cell 34: 1250–1272.

Gady, A, L, F., Vriezen, W, H., Van de Wal, M, H, B, J., Huang, P, P., Bovy, A, G., Visser, R. G, F., and Bachem, C, W, B. (2012). Induced point mutations in the phytoene synthase 1 gene cause differences in carotenoid content during tomato fruit ripening. Mol Breeding 29: 801–812.

Fujisawa, M., Nakano, T., Shima, Y., and Ito, Y. (2013). A large-scale identification of direct targets of the tomato MADS box transcription factor RIPENING INHIBITOR reveals the regulation of fruit ripening. The Plant Cell 25: 371–386.

Guo, J, E., Hu, Z, L., Zhu, M, K., Li, F, F., Zhu, Z, G., Lu, Y., and Chen, G, P. (2017). The tomato histone deacetylase *SlHDA1* contributes to the repression of fruit ripening and carotenoid accumulation. Sci Rep 7: 7930.

Guo, J, E., Hu, Z, L., Yu, X, H., Li, A, Z., Li, F, F., Wang, Y, S., Tian, S, B., and Chen, G, P. (2018). A histone deacetylase gene, *SlHDA3*, acts as a negative regulator of fruit ripening and carotenoid accumulation. Plant Cell Rep 37: 125–135.

Guo, J, E. (2022). Histone deacetylase gene *SlHDT1* regulates tomato fruit ripening by affecting carotenoid accumulation and ethylene biosynthesis. Plant Sci 318: 111235.

Hörtensteiner, S. (2009). Stay-green regulates chlorophyll and chlorophyll-binding protein degradation during senescence. Trends Plant Sci 14: 155–162.

Ilahy, R., Siddiqui, M, W., Tlili, I., Montefusco, A., Piro, G., Hdider, Chafik., and Lenucci, M, S. (2018). When color really matters: horticultural performance and functional quality of high-lycopene tomatoes. Crit Rev Plant Sci 37: 15–53.

Jiang, G, X., Li, Z, W., Ding, X, C., Zhou, Y, J., Lai, H, M., Jiang, Y, M., Duan, X, W. (2023). WUSCHEL-related homeobox transcription factor *SlWOX13* regulates tomato fruit ripening. Plant Physiol, kiad623. 10.1093/plphys/kiad623.

Jumper, J., Evans, R., Pritzel, A., Green, T., Figurnov, M., Ronneberger, O., Tunyasuvunakool, K., Bates, R., Žídek, A., Potapenko, A., Bridgland, A., Meyer, C., Kohl, S, A, A., Ballard, A, J., Cowie, A., Romera-Paredes, B., Nikolov, S., Jain, R., Adler, J., Back, T., Petersen, S., Reiman, D., Clancy, E., Zielinski, M., Steinegger, M., Pacholska, M., Berghammer, T., Bodenstein, S., Silver, D., Vinyals, O., Senior, A, W., Kavukcuoglu, K., Kohli, P., and Hassabis, D. (2021). Highly accurate protein structure prediction with AlphaFold. Nature 596: 583–589.

Kachanovsky, D, E., Filler, S., Isaacson, T., and Hirschberg, J. (2012). Epistasis in tomato color mutations involves regulation of phytoene synthase 1 expression by cis-carotenoids. P Natl Acad Sci USA 109: 19021–19026.

Kim, K, C., Lai, Z, B., Fan, B, F., and Chen, Z, X. (2008). *Arabidopsis WRKY38* and *WRKY62* transcription factors interact with histone deacetylase 19 in basal defense. Plant Cell 20: 2357–2371.

Kuang, J, F., Chen, J, Y., Luo, M., Wu, K, Q., Sun, W., Jiang, Y, M., and Lu, W, J. (2012). Histone deacetylase HD2 interacts with ERF1 and is involved in longan fruit senescence. J Exp Bot 63: 441–454.

Li, Z,W., Jiang, G, X., Liu, X, C., Ding, X, C., Zhang, D, D., Wang, X, W., Zhou, Y, J., Yan, H, L., Li, T, T., Wu, K, Q., Jiang, Y, M., and Duan, X, W. (2020). Histone demethylase *SlJMJ6* promotesfruit ripening by removing *H3K27* methylation of ripening-relatedgenes in tomato. New Phytol. 227:1138–1156.

Li, H., Wu, H., Qi, Q., Li, H, H., Li, Z, F., Chen, S., Ding, Q, Q., Wang, Q, Z., Yan, Z, M., Gai, Y., Jiang, X, N., Ding, J., Gu, T, T., Hou, X, L., Richard, M., Zhao, Y, D., and Li, Y. (2019). Gibberellins play a role in regulating tomato fruit ripening. Plant Cell physiol 60: 1619–1629.

Liang, Y, F., Bai, J, Y., Xie, Z, L., Lian, Z, Y., Guo, J., Zhao, F, Y., Liang, Y., Huo, H, Q., and Gong, H, J. (2023). Tomato sucrose transporter *SlSUT4* participates in flowering regulation by modulating gibberellin biosynthesis. Plant Physiol 192: 1080–1098.

Lin, T., Zhu, G, T., Zhang, J, H., Xu, X, Y., Yu, Q, H., Zheng, Z., Zhang, Z, H., Lun, Y, Y., Li, S., Wang, X, X., Huang, Z, J., Li, J, M., Zhang, C, H., Wang, T, T., Zhang, Y, Y., Wang, A, X., Zhang, Y, C., Lin, K., Li, C, Y., Xiong, G, S., Xue, Y, B., Mazzucato, A., Causse, M., Fei, Z, J., Giovannoni, J, J., Chetelat, R, T., Zamir, D., Städler, T., Li, J, F., Ye, Z, B., Du, Y, C., and Huang, S, W. (2014). Genomic analyses provide insights into the history of tomato breeding. Nat Genet 46: 1220–1226.

Liu, X, X., Huang, Y, G, M., Qiu, Z, K., and Gong, H. (2020). Comparative transcriptome analysis of differentially expressed genes between the fruit peel and flesh of the purple tomato cultivar ‘Indigo Rose’. Plant Signal Behav 15: 1752534.

Livak, K, J., and Schmittgen, T, D. (2001). Analysis of relative gene expression data using real-time quantitative PCR and the 2^−ΔΔCT^ method. Methods 25: 402–408.

Luo, Z, D., Zhang, J, H., Li, J, H., Yang, C, X., Wang, T, T., Ouyang, B., Li, H, X., Giovannoni, J., and Ye, Z, B. (2013). A STAY-GREEN protein SlSGR1 regulates lycopene and β-carotene accumulation by interacting directly with SlPSY1 during ripening processes in tomato. New Phytol 198: 442–452.

Martel, C., Vrebalov, J., Tafelmeyer, P., and Giovannoni, J, J. (2011). The tomato MADS-box transcription factor RIPENING INHIBITOR interacts with promoters involved in numerous ripening processes in a COLORLESS NONRIPENING-dependent manner. Plant Physiol 157: 1568–1579.

Ni, X., Ni, Z., Ouma, K, O., and Gao, Z, H. (2022). Mutations in *PmUFGT3* contribute to color variation of fruit skin in Japanese apricot (*Prunus mume* Sieb. et Zucc.). BMC Plant Biol 22: 1–16.

Petry, F, C., and Mercadante, A, Z. (2018). New method for carotenoid extraction and analysis by HPLC-DADMS/MS in freeze-dried Citrus and Mango pulps. J Brazil Chem Soc 1: 205–215.

Summers, C., Karst, F., and Charles, A, D. (1993). Cloning, expression and characterisation of the cDNA encoding human hepatic squalene synthase, and its relationship to phytoene synthase. Gene 136: 185–192.

Sun, T, H., Rao, S, B., Zhou, X, S., and Li, L. (2022). Plant carotenoids: Recent advances and future perspectives. Mol Hortic 2: 3.

Wang, L., Zhang, X, L., Wang, L., Tian, Y, N., Jia, N., Chen, S, Z., Shi, N, B., Huang, X, M., Zhou, C., Yu, Y, W., Zhang, Z, Q., and Pang, X, Q. (2017). Regulation of ethylene-respons ive *SlWRKYs* involved in color change during tomato fruit ripening. Sci Rep 7: 16674.

Wang, W, H., Wang, P, W., Li, X, J., Wang, Y, Y., Tian, S, P., and Qin, G, Z. (2021). The transcription factor *SlHY5* regulates the ripening of tomato fruit at both the transcriptional and translational levels. Hortic Res 8: 83.

Wang, Z, R., Gao, M., Li, Y, F., Zhang, J, L., Su, H., Cao, M., Liu, Z, J., Zhang, X, C., Zhao, B., Guo, Y, D., and Zhang, N. (2022). The transcription factor *SlWRKY37* positively regulates jasmonic acid-and dark-induced leaf senescence in tomato. J Exp Bot 73: 6207–6225.

Wu, M, B., Xu, X., Hu, X, W., Liu, Y, D., Cao, H, H., Chan, H, L., Gong, Z, H., Yuan, Y, J., Luo, Y, Q., Feng, B, H., Li, Z, G., and Deng, W. (2020). *SlMYB72* regulates the metabolism of chlorophylls, carotenoids, and flavonoids in tomato fruit. Plant Physiol 183: 854–868.

Xiong, C., Luo, D., Lin, A, H., Zhang, C, L., Shan, L, B., He, P., Li, B., Zhang, Q, M., Hua, B., Yuan, Z, L., Li, H, X., Zhang, J, H., Yang, C, X., Lu, Y, G., Ye, Z, B., and Wang, T, T. (2019). A tomato B-box protein *SlBBX20* modulates carotenoid biosynthesis by directly activating PHYTOENE SYNTHASE 1, and is targeted for 26S proteasome-mediated degradation. New Phytol 221: 279–294.

Xu, Z, J., Wang, J,C., Ma, Y, B., Wang, F., Wang, J, R., Zhang, Y., and Hu, X, H. (2023). The bZIP transcription factor *SlAREB1* regulates anthocyanin biosynthesis in response to low temperature in tomato. Plant J 115: 205–219.

Yang, X, T., Pang, X, Q., Xu, L, Y., Fang, R, Q., Huang, X, M., Guan, P, J., Lu, W, J., and Zhang, Z, Q. (2009). Accumulation of soluble sugars in peel at high temperature leads to stay-green ripe banana fruit. J Exp Bot 60: 4051–4062.

Yuan, H., Owsiany, K., Sheeja, T, E., Zhou, X, J., Rodriguez, C., Li, Y, X., Welsch, R., Chayut, N., Yang, Y., Thannhauser, T, W., Parthasarathy, M, V., Xu, Q., Deng, X, X., Fei, Z, J., Schaffer, A., Katzir, N., Burger, J., Tadmor, Y., and Li, L. (2015). A single amino acid substitution in an ORANGE protein promotes carotenoid overaccumulation in *Arabidopsis*. Plant Physiol 169: 421–431.

Yuan, Y., Ren, S, Y., Liu, X, F., Su, L, Y., Wu, Y., Zhang, W., Li, Y., Jiang, Y, D., Wang, H, H., Fu, R., Bouzayen, M., Liu, M, C., and Zhang, Y. (2022). *SlWRKY35* positively regulates carotenoid biosynthesis by activating the MEP pathway in tomato fruit. New Phytol 234: 164–178.

Zhao, L, M., Lu, J, X., Zhang, J, X., Wu, P, Y., Yang, S, G., and Wu, K, Q. (2015). Identification and characterization of histone deacetylases in tomato (*Solanum lycopersicum*). Front Plant Sci 5: 760.

Zhang, J., Li, H., Huang, X, R., Xing, J., Yao, J, M., Yin, T, C., Jiang, J, F., Wang, P, C., and Xu, B. (2022). STAYGREEN-mediated chlorophyll a catabolism is critical for photosystem stability during heat-induced leaf senescence in perennial ryegrass. Plant Cell Environ 45: 1412–1427.

Zhang, Y., Yang, X., Nvsvrot, T., Huang, L, Y., Cai, G, H., Ding, Y, W., Ren, W, Y., and Wang, N. (2022). The transcription factor *WRKY75* regulates the development of adventitious roots, lateral buds and callus by modulating hydrogen peroxide content in poplar. J Exp Bot 73: 1483–1498.

Zhang, L, L., Zheng, Y., Xiong, X, X., Li, H., Zhang, X., Song, Y, L., Zhang, X, H., and Min, D, H. (2023). The wheat VQ motif-containing protein TaVQ4-D positively regulates drought tolerance in transgenic plants. J Exp Bot 74: 5591–5605.

